# CEACAM5/6^+^ Tumor Cells and IL-1β^+^ Macrophages Drive Resistance to Chemo-immunotherapy in Gastric Cancer

**DOI:** 10.64898/2026.03.05.708917

**Authors:** Jian Chen, Liudeng Zhang, Yikai Luo, Xiaying Han, Muxing Kang, Jing Chen, Wei Liu, Zhenzhen Xun, Guofeng Chen, Ke Chen, Shenbin Xu, Chaoyang Zhang, Zhiwei Wu, Wenxuan Wu, Zhixing Hao, Yaxuan Han, Qiaowei Lin, Yewei Xu, Lie Wang, Han Liang

**Affiliations:** Department of Gastrointestinal Surgery, The Second Affiliated Hospital, Zhejiang University School of Medicine, Hangzhou 310000, China; Cancer Center, Zhejiang University, Hangzhou 310058, China; Graduate Program in Quantitative and Computational Biosciences, Baylor College of Medicine, Houston, TX 77030, USA; Department of Bioinformatics and Computational Biology, The University of Texas MD Anderson Cancer Center, Houston, TX 77030, USA; Institute of Immunology and Bone Marrow Transplantation Center, The First Affiliated Hospital, Zhejiang University School of Medicine, Hangzhou 310006, China; Liangzhu Laboratory, Zhejiang University Medical Center, Hangzhou 311121, China; Department of Systems Biology, The University of Texas MD Anderson Cancer Center, Houston, TX 77030, USA

## Abstract

Chemo-immunotherapy is a first-line treatment for advanced gastric cancer, yet response rates remain limited and resistance mechanisms are poorly defined. Here we generate a single-cell atlas of 542,121 cells from 35 patients treated with anti–PD-1 plus chemotherapy, profiling pre- and post-treatment tumors linked to clinical response. Integrating spatial transcriptomics, immunohistochemistry, and bulk RNA sequencing, we identify two temporally distinct resistance programs. Intrinsic resistance in pre-treatment non-responders is marked by enrichment of CEACAM5/6⁺ tumor cells that form immune-excluded spatial niches characterized by macrophage recruitment and CD8⁺ T-cell exhaustion. Acquired resistance in post-treatment non-responders is driven by expansion of IL-1β⁺ macrophages, which induces coordinated NF-κB activation across tumor and stromal compartments, promoting PD-L1 upregulation, epithelial–mesenchymal transition, and chronic inflammation. These findings delineate an evolutionary trajectory of resistance and nominate CEACAM5/6 and IL-1β as predictive biomarkers and therapeutic targets to improve anti–PD-1–based combination strategies.

## Introduction

Immune checkpoint blockade has transformed cancer treatment, yet in gastric cancer, most patients either fail to respond or develop resistance within months of initial response (1,2). The CheckMate-649 and KEYNOTE-859 trials established anti-PD-1 antibodies combined with chemotherapy as the standard of care for advanced gastric cancer, but objective response rates remain limited, ranging from 40-60% even in biomarker-selected populations, and the majority of responding patients eventually progress (3,4). Resistance to immune checkpoint blockade arises through diverse mechanisms, including tumor-intrinsic immune evasion, loss of antigen presentation, and immunosuppressive remodeling of the tumor microenvironment (TME) (5–9). In gastric cancer, the interplay between tumor cell-intrinsic programs and the immunosuppressive TME remains poorly understood, limiting the rational design of combination strategies. This clinical reality—dramatic responses in some patients, rapid failure in others—underscores the urgent need to identify molecular features that predict treatment outcome or define new clinically actionable targets.

Single-cell RNA sequencing has begun to illuminate the cellular heterogeneity underlying immunotherapy response in gastric cancer (10–14), identifying candidate resistance-associated populations such as NKG2A^+^ T cells (15), CLEVER-1^+^ macrophages (16), inflammatory cancer-associated fibroblasts, and SPP1^+^ tumor-associated macrophages (17,18). However, the question of whether resistance-associated features are present before treatment initiation or emerge during therapy remains incompletely resolved. Systematic comparison of pre- and post-treatment tumors stratified by clinical response can distinguish features associated with intrinsic versus acquired resistance — a distinction with direct clinical implications, as biomarkers for upfront patient selection differ from those guiding intervention upon disease progression.

Here, we present a large-scale single-cell RNA sequencing atlas comprising 542,121 cells from 35 gastric cancer patients treated with anti-PD-1–based chemo-immunotherapy, with tumor samples collected prior to treatment initiation or following chemo-immunotherapy and linked to clinical response. This study design enables systematic comparison of responders and non-responders across distinct treatment phases. By delineating the molecular and cellular programs associated with both intrinsic and acquired resistance, our findings establish a comprehensive framework for understanding therapeutic failure and improving chemo-immunotherapy in gastric cancer.

## Results

### A single-cell atlas of gastric cancer under anti-PD-1-based chemo-immunotherapy

To systematically characterize resistance mechanisms to anti-PD-1-based chemo-immunotherapy in gastric cancer, we performed single-cell RNA sequencing on tumor samples from 35 patients receiving anti-PD-1 antibodies in combination with chemotherapy (**Fig. 1A, Table S1**). Patients were stratified into responders (R) and non-responders (NR) based on Response Evaluation Criteria in Solid Tumors (RECIST). Samples were collected at two timepoints: before treatment initiation and during surgical resection following therapy. Our cohort comprised 70 clinical specimens collected from diverse anatomical sites, including primary gastric tumors, liver metastases, ovarian metastases, peripheral blood, and lymph nodes. Given that the primary tumor site represents the major disease burden and therapeutic target, we focused our analyses on these gastric samples (n = 32; 20 pre-treatment, 12 post-treatment), of which 8 pre-treatment and 11 post-treatment samples had confirmed response classifications following neoadjuvant chemo-immunotherapy. This rich dataset enables four key comparisons: pre-treatment R versus NR, post-treatment R versus NR, pre-versus post-treatment within responders, and pre-versus post-treatment within non-responders (**Fig. 1A**). To validate findings from this cohort, we performed histopathological confirmation using H&E staining and extended our analysis across published datasets, including single-cell RNA sequencing, bulk RNA sequencing, and spatial transcriptomics (**Table S2**).

**Figure 1.**
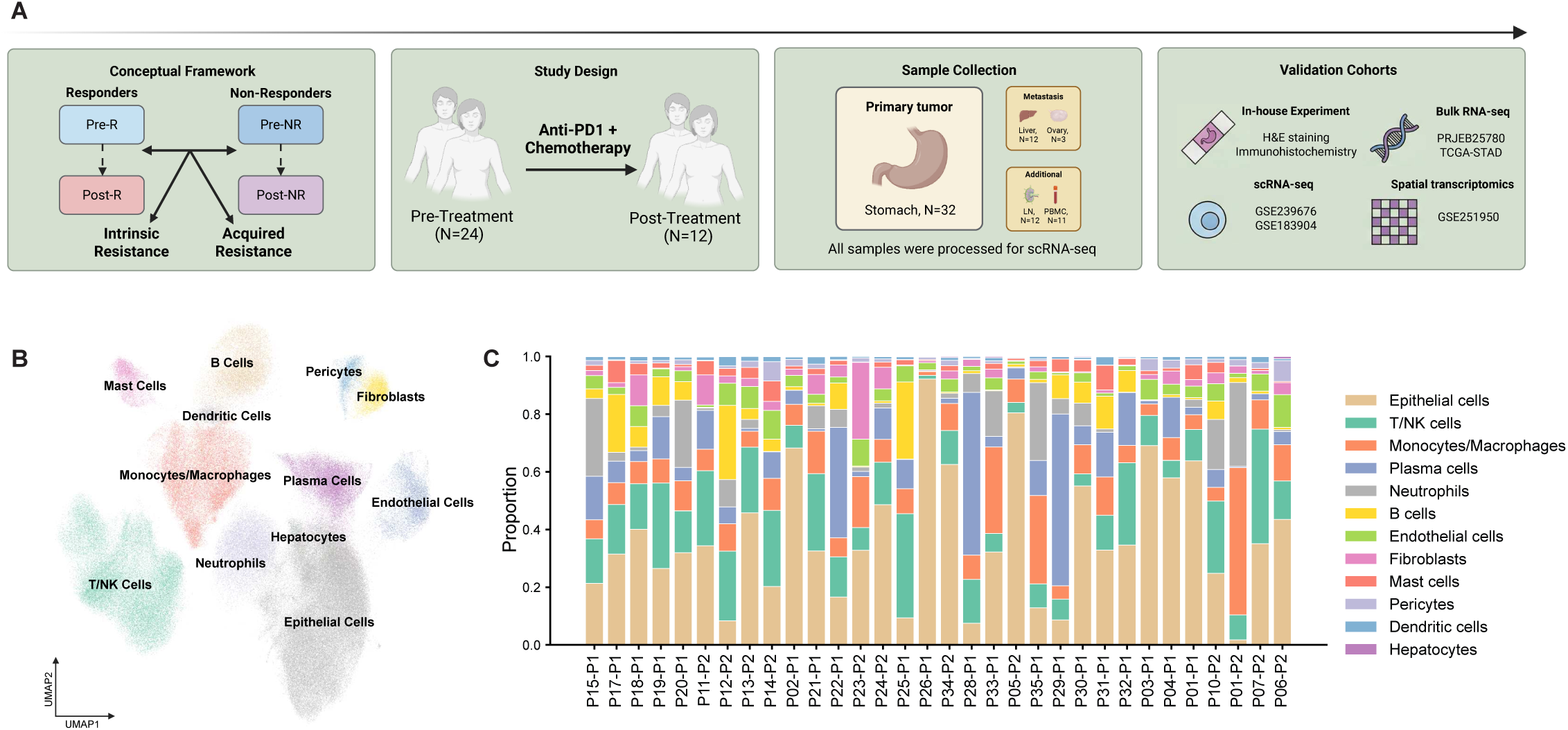
A single-cell transcriptomic atlas of gastric cancer under anti-PD-1-based chemo-immunotherapy. (A) Schematic of study design. (B) UMAP embedding of 542,121 cells colored by 12 major cell types. (C) Proportion of each major cell type per sample (n = 32).

After rigorous quality control and doublet removal, we obtained 542,121 high-quality single cells for downstream analysis (**Fig. S1A–E**). Unsupervised clustering identified 12 major cell clusters. The T/NK cluster was further resolved into CD4^+^ T cells, CD8^+^ T cells, and NK cells through sub-clustering, yielding 14 annotated cell types: epithelial cells, CD4^+^ T cells, CD8^+^ T cells, NK cells, B cells, plasma cells, monocytes/macrophages, dendritic cells, neutrophils, fibroblasts, endothelial cells, mast cells, pericytes, and hepatocytes, visualized as distinct populations in uniform manifold approximation and projection (UMAP) (**Fig. 1B**). Cell type annotation was validated using canonical marker gene expression (**Fig. S1F,G**). Analysis of cell type distribution across the 32 gastric tumor samples revealed inter-patient heterogeneity in immune infiltration (**Fig. 1C**). Cell type compositions from metastatic sites, including liver metastases (n = 12; 10 pre-treatment, 2 post-treatment), are presented in **Fig. S1H**. This comprehensive atlas comprising 14 major cell types and 84 minor cell states, with primary focus on gastric tumors, provided the foundation for investigating both tumor-intrinsic and TME-mediated resistance mechanisms.

### CEACAM5/6^+^ tumor cells are enriched in pre-treatment non-responders

Resistance to immune checkpoint blockade can arise from tumor-intrinsic mechanisms present before treatment initiation (5–9). To identify such pre-existing features, we analyzed pre-treatment epithelial cells, reasoning that molecular differences between responders and non-responders at baseline would reflect intrinsic resistance determinants and potential therapeutic targets. These cells are characterized by elevated copy-number variation, a hallmark feature of malignant tumor cells. Unbiased Leiden clustering of pre-treatment epithelial cells resolved nine transcriptionally distinct subpopulations/cell states (**Fig. 2A, Fig. S2A**). Among them, one cluster was distinguished by markedly elevated expression of both CEACAM5 and CEACAM6 (**Fig. 2B, C**). These two genes exhibited a strong positive correlation (Spearman ρ = 0.93, *P* < 1×10⁻¹⁶; *n* = 49,696 pre-treatment epithelial cells; **Fig. 2D**), a finding further validated in two independent datasets (**Fig. S2B, C**). This coordinated transcriptional pattern led us to designate this subpopulation as Epi_CEACAM5/6⁺. CEACAM5 and CEACAM6 are frequently overexpressed in human cancers, with CEACAM5 detected in up to 70% of solid tumors (19,20). Functionally, CEACAM5 promotes immune evasion through interactions with inhibitory receptors on immune cells, while CEACAM6 directly suppresses cytotoxic T cell responses via the CEACAM6–CEACAM1 axis; both are highly expressed in cancer types resistant to PD-1/PD-L1 blockade (19,21–24).

**Figure 2.**
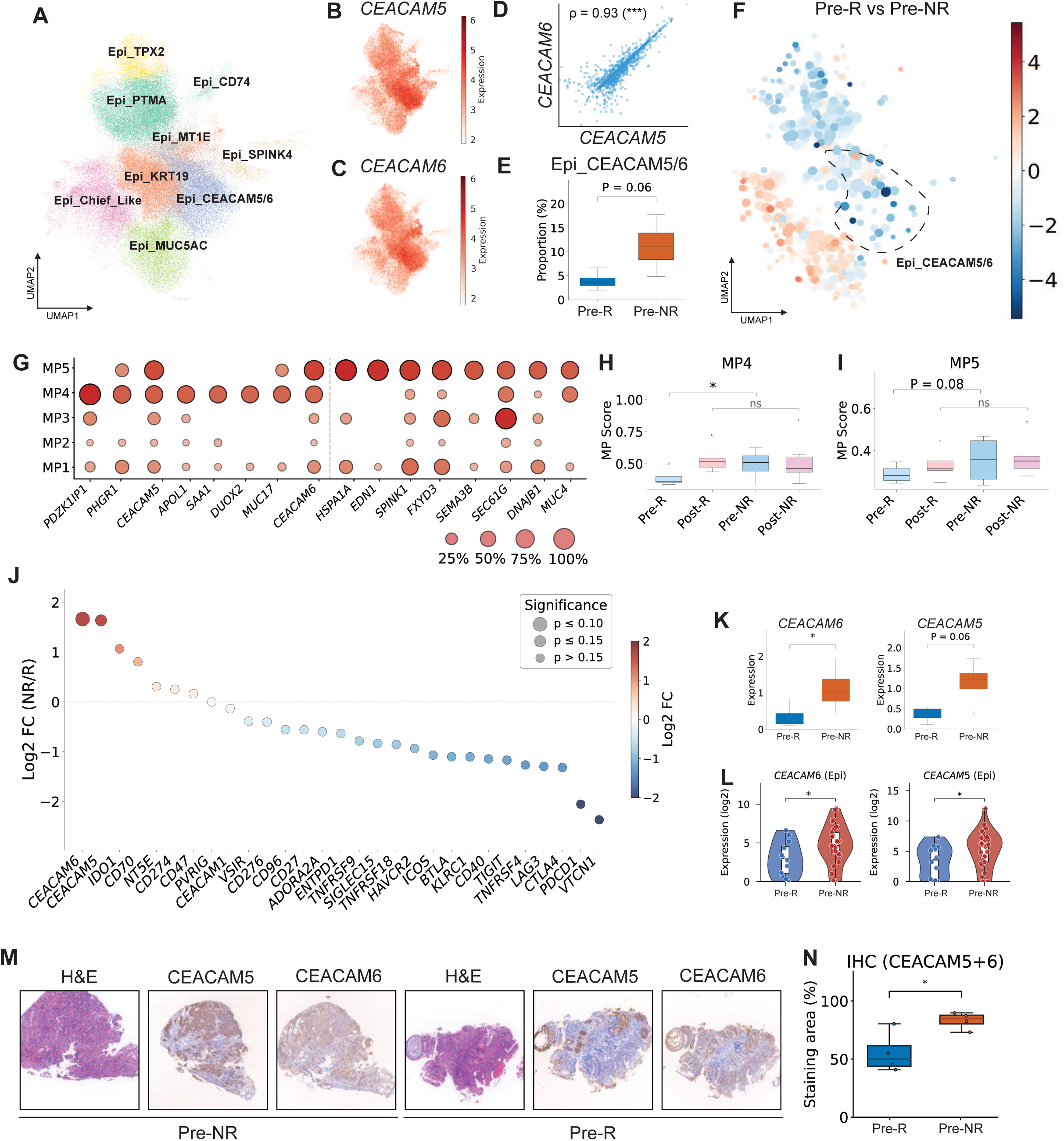
Enrichment of CEACAM5/6^+^ tumor cells in pre-treatment non-responder tumors. (A) UMAP of epithelial cells colored by 9 cell states. (B, C) UMAP colored by CEACAM5 (B) and CEACAM6 expression (log-normalized). (D) Spearman rank correlation between CEACAM5 and CEACAM6 in pre-treatment epithelial cells. (E) Proportion of CEACAM5/6^+^ epithelial cells per sample in pre-treatment R (n = 4) versus NR (n = 4). P-value, Mann–Whitney U test. (F) Milo differential abundance analysis of epithelial neighborhoods between pre-treatment R and NR samples. Color indicates log-fold change (red, NR-enriched; blue, R-enriched); size indicates neighborhood cell count. (G) Gene loading dotplot for top genes per meta-program. (H, I) MP4 (H) and MP5 (I) activity scores across different treatment–response groups. Exact permutation test (Post-R vs. all others) and Kruskal–Wallis test. (J) Pseudo-bulk epithelial-cell expression levels of immune checkpoint genes with clinical trials comparing pre-treatment R versus NR. Dot size, significance; color, log_2_ fold change. P-values, Mann–Whitney U test. (K) Sample-level CEACAM5 and CEACAM6 expression in pre-treatment R versus NR. One-tailed Mann–Whitney U test. (L) Validation of CEACAM5 and CEACAM6 expression in an independent cohort under anti-PD1 treatment (PRJEB25780; BayesPrism-deconvolved epithelial expression). One-tailed Mann–Whitney U test. (M) Representative H&E, CEACAM5, and CEACAM6 immunohistochemistry (IHC) images from pretreatment non-responder (NR) and responder (R) tumors. (N) Combined CEACAM5 + CEACAM6 IHC staining area in R (n = 4) versus NR (n = 4). One-tailed Mann–Whitney U test. Each dot represents one patient. *, *P* < 0.05; ***, *P* < 10^-3^.

Sample-level quantification revealed that the CEACAM5/6^+^ cluster was enriched in non-responders (11.2% ± 4.7% vs. 4.0% ± 1.7% in responders, 2.8-fold, P = 0.06, exact permutation test; **Fig. 2E**); A similar trend was observed in liver metastasis, though it did not reach statistical significance (**Fig. S2D**). To confirm that this enrichment was not an artifact of discrete cluster assignment, we performed differential abundance analysis using Milo—an unsupervised method that tests for compositional shifts across overlapping cell neighborhoods on the KNN graph without requiring discrete cluster boundaries (51). Milo identified the CEACAM5/6^+^ region as significantly enriched in non-responders (SpatialFDR < 0.1; **Fig. 2F**), and the neighborhoods enriched in non-responders closely overlapped with regions of high CEACAM5/6 expression in the UMAP embedding, providing independent confirmation of the Leiden-based finding.

We next asked whether this enrichment was also evident at the level of transcriptional programs. An orthogonal analysis using per-sample non-negative matrix factorization (NMF) identified five metaprograms (25), of which MP4 (defined by mucin family members MUC5AC, MUC5B, MUC6 and secretory markers) and MP5 (characterized by proliferation-associated genes) showed the strongest association with CEACAM5/6 (**Fig. 2G, Fig. S2EF, Table S3**). Both CEACAM5 and CEACAM6 ranked among the top-weighted genes in MP4 and MP5. Consistent with the cell-state based analysis, MP4 was significantly depleted in pre-treatment non-responders comparing to other samples (P = 0.03, exact permutation test), and MP5 showed the same trend with marginal significance (P = 0.08; **Fig. 2H, I**), linking CEACAM5/6-driven transcriptional programs with treatment outcome at the functional level.

To further confirm the robustness of our findings, we performed pseudo-bulk differential expression analysis of immune checkpoint and immunomodulatory receptors with approved drugs or clinical trials (**Table S4**). Among all actionable targets, CEACAM5 and CEACAM6 emerged as among the most significantly upregulated in non-responders (**Fig. 2J**). Sample-level quantification revealed significantly elevated CEACAM6 expression in non-responders (mean 1.11 vs. 0.35, 3.2-fold, P < 0.05, exact permutation test) and the same trend with marginal significance for CEACAM5 (mean 1.14 vs. 0.37, 3.1-fold, P = 0.06, exact permutation test; **Fig. 2K**), reinforcing CEACAM5/6 as a defining feature of the tumor resistance phenotype. These findings were independently validated in another gastric cancer immunotherapy cohort (PRJEB25780; n = 12 responders, n = 33 non-responders) (26), where non-responders showed significantly elevated CEACAM5 and CEACAM6 expression (CEACAM5, P = 0.026, CEACAM6, P = 0.03, Mann–Whitney U test **Fig. 2L**). At the protein level, we performed immunohistochemical analysis on independent samples (**Table S5**) and confirmed significantly higher combined CEACAM5/6 staining in non-responder tumors (83.2% vs. 55.3%, P = 0.03, Mann–Whitney U test, n = 4 per group; **Fig. 2M, N**).

Together, these convergent lines of evidence—unsupervised clustering, differential abundance testing, NMF-based functional programs, pseudo-bulk differential expression, external cohort validation, and protein-level confirmation by immunohistochemistry—establish CEACAM5/6^+^ tumor cells as a robust marker of intrinsic resistance to anti-PD-1 therapy in gastric cancer (notably, CEACAM5/6 expression was not associated with patient survival, suggesting that it is not a prognostic marker, **Fig. S2G**). The consistency across multiple analytical approaches and independent datasets underscores the potential of CEACAM5/6 as both a predictive biomarker and a therapeutic target.

### CEACAM5/6 expression correlates with CD8^+^ T cell exhaustion and the establishment of an immunosuppressive spatial niche

Having established that CEACAM5/6^+^ tumor cells are enriched in pre-treatment non-responders, we next investigated whether CEACAM5/6 expression is associated with broader changes in the TME, focusing on immune checkpoint ligand expression, CD8^+^ T cell exhaustion, and spatial organization. At the sample level, CEACAM5 and CEACAM6 expression in epithelial cells demonstrated significant or marginally significant positive correlations with PD-L1 expression (**Fig. 3A**), linking CEACAM5/6 to immune checkpoint signaling. These associations were independently validated in TCGA gastric cancer samples using deconvolved epithelial expression (**Fig. 3B**), confirming a robust connection between CEACAM5/6 and PD-L1 upregulation.

**Figure 3.**
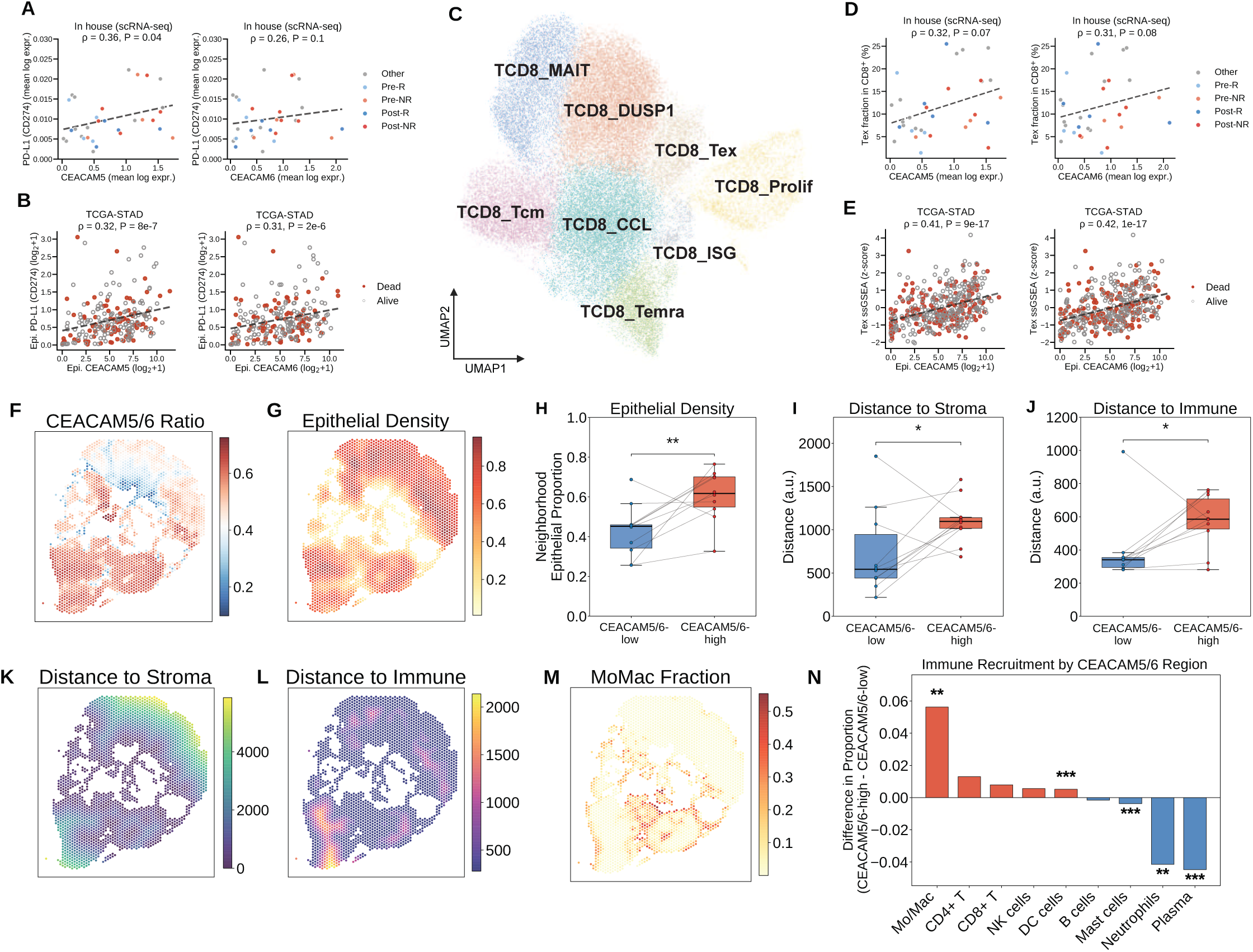
CEACAM5/6 expression correlates with CD8^+^ T cell exhaustion and shapes the spatial immune microenvironment. (A) Spearman rank correlation between sample-level CEACAM5 (left) or CEACAM6 (right) and CD274 (PD-L1) expression in pre-treatment epithelial cells (n = 31 samples). Points colored by treatment–response group. (B) Spearman rank correlation between CEACAM5 (left) or CEACAM6 (right) and CD274 in TCGA-STAD (n = 407; BayesPrism-deconvolved). (C) UMAP of CD8^+^ T cells colored by 8 cell states. (D) Same as (A) for CD8^+^ exhausted T cell (Tex) fraction. (E) Same as (B) for Tex ssGSEA score (18-gene signature; see Methods). (F, G) Spatial maps of a representative sample showing CEACAM5/6^+^ expression ratio (F) and epithelial neighborhood density (G). (H–J) Paired comparisons between CEACAM5/6-high and CEACAM5/6-low regions for epithelial neighborhood proportion (H), distance to nearest stromal region (I), and distance to nearest immune region (J). One-tailed Wilcoxon signed-rank test (paired). (K, L) Spatial maps of distance to stromal boundary (K) and immune region (L). (M) Spatial map of deconvolved monocyte/macrophage fraction. (N) Difference in immune cell type proportions between CEACAM5/6-high and CEACAM5/6-low regions. Red, enriched in CEACAM5/6-high; blue, enriched in CEACAM5/6-low. *, *P* < 0.05; **, *P* < 0.01; ***, *P* < 10^-3^.

CEACAM5/6 expression has been reported to link with CD8^+^ T cell exhaustion (24). Among 8 transcriptionally distinct CD8^+^ T cell states identified by sub-clustering (**Fig. 3C**), one cluster was annotated as exhausted based on elevated expression of canonical markers PDCD1, HAVCR2 and a composite Tex score (**Fig. S3**). Sample-level Tex proportion correlated positively with CEACAM5 and CEACAM6 expression (**Fig. 3D**), stronger associations reproduced in the TCGA gastric cancer cohort (**Fig. 3E**). Together with the PD-L1 correlations, these findings suggest a mechanistic link in which CEACAM5/6 promotes PD-L1 upregulation on tumor cells, which in turn drives CD8^+^ T cell exhaustion through sustained PD-1/PD-L1 engagement—consistent with CEACAM5/6 acting as an upstream correlate of both checkpoint ligand expression and adaptive immune dysfunction.

We next asked whether CEACAM5/6^+^ cells also shape the spatial organization of the TME to promote immune exclusion. CEACAM family members have been implicated in intercellular adhesion and tissue architecture (19,20), and their overexpression may physically reorganize the tumor–stroma interface, creating barriers to immune cell infiltration. Spatial transcriptomics data from an independent gastric cancer cohort (GSE251950) (27–29) revealed marked heterogeneity in CEACAM5/6 expression across tumor regions, with CEACAM-high epithelium preferentially localized to areas of high tumor cell density (**Fig. 3F, G**). These spatial patterns were observed across all 10 samples in the cohort (**Fig. S4**). CEACAM-high regions exhibited significantly elevated neighborhood epithelial density compared to CEACAM-low regions within the same tumors (P = 0.01, Wilcoxon signed-rank test; **Fig. 3H**), indicating that CEACAM5/6 expression marks densely packed tumor cores. This is consistent with the known role of CEACAM5/6 in homophilic cell–cell adhesion, which promotes tight epithelial packing and may limit immune cell infiltration into the tumor parenchyma (19,21).

Distance analysis further showed that CEACAM5/6-high regions were significantly further from stromal cells (P = 0.04, Wilcoxon signed-rank test; **Fig. 3I**) and immune-rich areas (P = 0.03; **Fig. 3J**), consistent with an immune exclusion phenotype. Spatial distance mapping confirmed increased separation of CEACAM5/6-high regions from both stromal (**Fig. 3K**) and immune (**Fig. 3L**) populations. A linear mixed-effects model adjusting for tumor density revealed that CEACAM5/6-high epithelium was significantly associated with increased recruitment of monocytes/macrophages and neutrophils (positive coefficients, P < 10^-3^; see Methods) but showed negative or null associations with adaptive immune populations including T cells, B cells, and NK cells. Spatial deconvolution corroborated these findings, showing selective enrichment of monocytes/macrophages in CEACAM5/6-high areas (**Fig. 3M**), while region-level analysis demonstrated that CEACAM5/6-high dominant tumor regions were enriched for macrophages and neutrophils but depleted of CD4^+^ T cells, CD8^+^ T cells, and B cells (**Fig. 3N**). This pattern of selective myeloid recruitment with adaptive immune exclusion is consistent with the broader literature on myeloid-mediated immunosuppression in solid tumors (30–34).

Together, these analyses support a model in which CEACAM5/6^+^ cancer cells are associated with an immunosuppressive spatial niche characterized by high tumor density, selective recruitment of myeloid cells, and exclusion of adaptive immune populations—a spatial organization that likely limits the efficacy of anti-PD-1 therapy by preventing T cell access to tumor cells.

### IL-1β+ macrophages selectively expand in post-treatment non-responders

Having characterized CEACAM5/6 as a tumor-intrinsic resistance mechanism in pre-treatment tumors, we next asked whether the immune compartment undergoes resistance-associated remodeling during treatment. Rather than focusing on individual cell types a priori, we adopted a systematic, unbiased approach: sample-level proportions for all immune cell states were computed across the 32 stomach samples, pairwise Spearman correlations identified co-varying cell states (**Fig. 4A**), and hierarchical clustering grouped co-regulated states into five distinct immune modules (IM), each enriched for specific immune cell populations: IM-T/NK/DC, IM-MoMac, IM-Mixed, IM-Neutrophil, and IM-B/Plasma (**Fig. 4B**).

**Figure 4.**
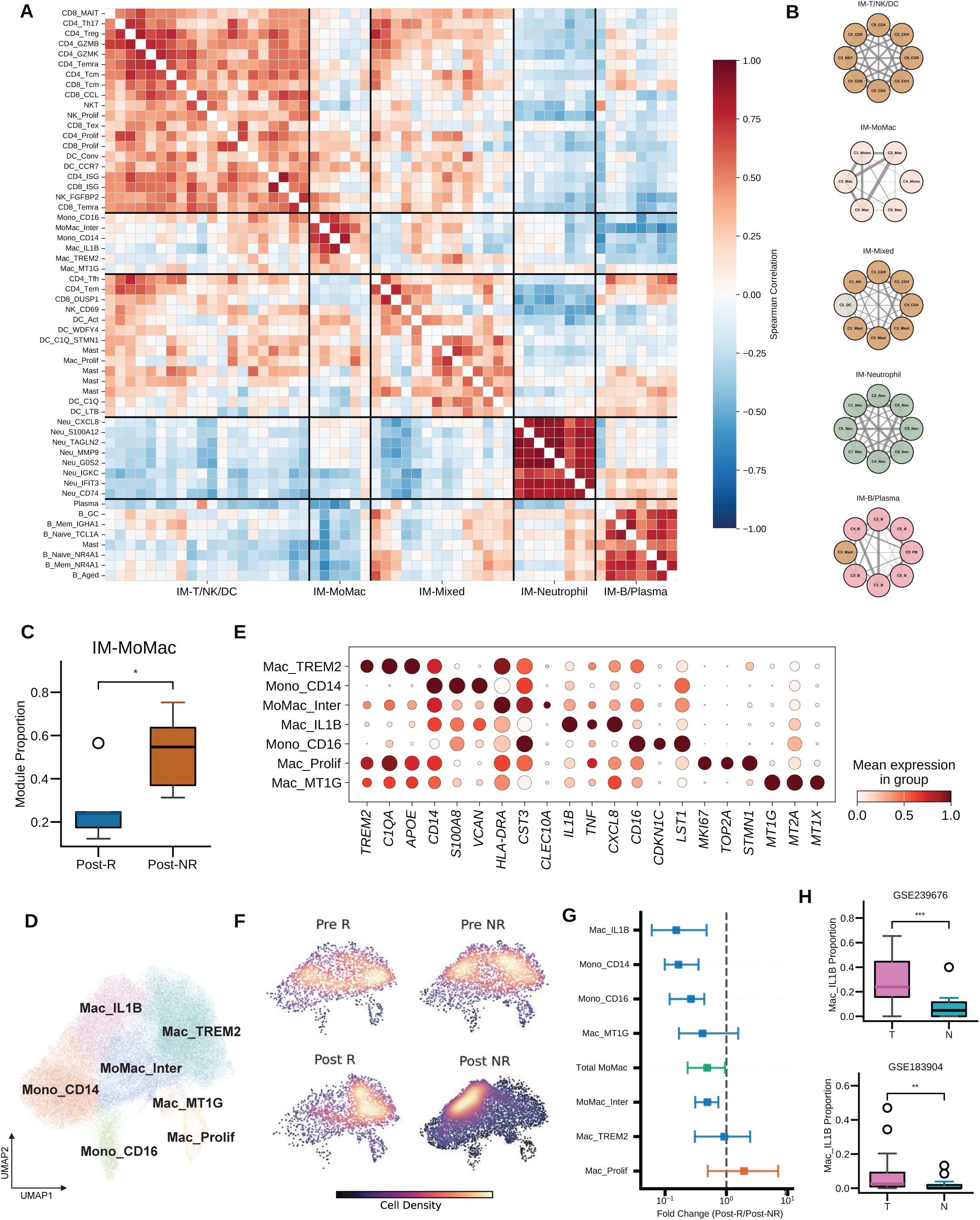
Enrichment of IL-1β^+^ macrophages correlates with acquired resistance. (A) Pearson correlation heatmap of immune cell state proportions across all stomach samples, organized by five immune modules (M1–M5) identified by hierarchical clustering. (B) Jaccard similarity network graphs for each module, showing cell state co-occurrence. Node color indicates cell type; edge thickness proportional to Jaccard similarity. (C) Module 2 (Monocyte/Macrophage module) proportion between post-treatment R (n = 5) and NR (n = 6) samples. P value, Mann–Whitney U test. (D) UMAP of Monocytes/Macrophages colored by 7 cell states. (E) Dotplot of canonical marker genes across 7 MoMac cell states. Dot size, percentage of expressing cells; color, mean expression. (F) Gaussian KDE of MoMac cell distribution across UMAP space, stratified by treatment phase and response. (G) Forest plot of fold changes for each MoMac cell state’s association with non-response in post-treatment samples. Blue, R-associated; red, NR-associated. (H) External validation of Mac_IL1B proportion in tumor versus adjacent normal tissue from two independent gastric cancer scRNA-seq cohorts (GSE239676, top; GSE183904, bottom). P-value, Mann–Whitney U test. *, *P* < 0.05; **, *P* < 0.01; ***, *P* < 10^-3^.

Differential abundance analysis between post-treatment responders and non-responders identified the IM-MoMac module as significantly elevated in non-responders (P = 0.03, Mann–Whitney U test; **Fig. 4C**), while the remaining modules—including those comprising T/NK cells and B/plasma cells—showed no significant association with treatment response (**Fig. S5**), indicating that resistance-associated remodeling is selective to the monocyte/macrophage compartment. Notably, this module comprised almost exclusively monocyte/macrophage subpopulations—a result that emerged from an unbiased analysis encompassing all immune cell types without prior selection (**Fig. 4 A, B**). This unbiased identification of macrophages as the dominant resistance-associated population aligns with emerging evidence that tumor-associated macrophages modulate therapeutic response across multiple cancer types (30–34), prompting us to investigate macrophage heterogeneity in greater detail.

Sub-clustering of the expanded monocyte/macrophage compartment resolved 7 distinct subpopulations with divergent transcriptional identities (**Fig. 4D**). Canonical marker gene expression defined these subpopulations (**Fig. 4E**), with IL-1β+ macrophages distinguished by high IL1B and TNF expression. Density analysis revealed that non-responders exhibited pronounced enrichment of specific macrophage subsets rather than uniform expansion, with the NR-enriched UMAP region corresponding to the IL-1β+ macrophage cluster (**Fig. 4F**). This heterogeneity is in line with the established plasticity of tumor-associated macrophages, which can adopt phenotypes ranging from anti-tumoral to pro-tumoral depending on microenvironmental cues (33,34).

To statistically validate this observation, we performed fold change analysis across all monocyte/macrophage subtypes (**Fig. 4G**). The IL-1β^+^ macrophage subset was confirmed as the most substantially enriched population in non-responders (fold change R/NR = 0.15, 95% CI [0.06, 0.47], P = 0.05, bootstrap), along with circulating monocytes. IL-1β-producing macrophages have been implicated in promoting tumor progression and therapy resistance through suppression of anti-tumor immunity and promotion of angiogenesis (35). Notably, other macrophage subsets showed protective associations (fold change > 1), underscoring that resistance is driven by a specific functional state rather than general macrophage accumulation. The enrichment of IL-1β^+^ macrophages in non-responders likely reflects an extension of their tumor-specific role. In two independent single-cell cohorts, IL-1β^+^ macrophages were markedly enriched in tumor tissues over adjacent normal tissues (P = 0.001, GSE239676; P = 0.01, GSE183904 (11); Mann–Whitney U test; **Fig. 4H**), consistent with active recruitment or local differentiation within the TME.

### IL-1β^+^ macrophages orchestrate pan-TME NF-κB activation and inflammatory reprogramming

The identification of IL-1β^+^ macrophages as the most strongly resistance-associated macrophage subtype was based on their differential abundance in non-responders. We next asked whether this enrichment was accompanied by functional alterations in the broader monocyte/macrophage compartment. Unbiased pathway enrichment analysis (GSEA) of all monocytes/macrophages revealed a coherent inflammatory signature in non-responders: TNFα signaling via NF-κB, inflammatory response, interferon gamma response, and IL-6/JAK/STAT3 signaling were among the most significantly enriched pathways (**Fig. 5A**). Critically, spatial mapping of these pathway scores across the macrophage UMAP embedding showed that all enriched pathways peaked within the IL-1β^+^ macrophage cluster region (**Fig. 5B**), demonstrating that the MoMac-module–level functional changes are driven by the IL-1β^+^ macrophage subpopulation rather than representing a compartment-wide phenomenon.

**Figure 5.**
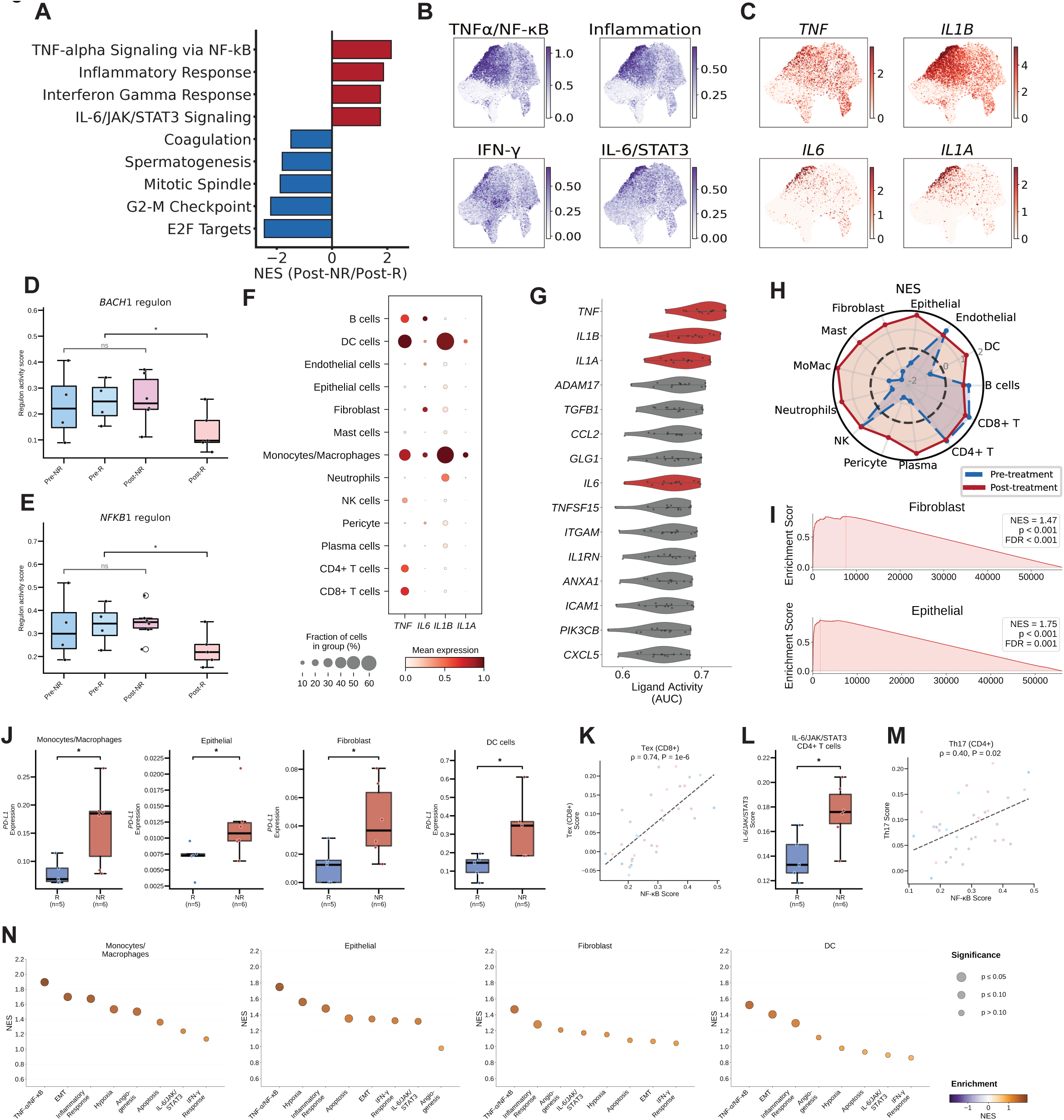
IL-1β^+^ macrophage-mediated NF-κB signaling drives immunosuppressive remodeling of the tumor microenvironment. (A) Top 9 Hallmark pathways enriched in Monocytes/Macrophages (post-treatment NR versus R), ranked by normalized enrichment score. GSEA pre-rank analysis. (B) UMAP of MoMac cells colored by pathway activity scores for TNF-α Signaling via NF-κB, Inflammatory Response, IFN-gamma Response, and IL-6/JAK/STAT3 Signaling. (C) UMAP of MoMac cells colored by TNF, IL1B, IL6, and IL1A expression. (D) BACH1 regulon activity in Mac_IL1B cells across four treatment–response groups. Exact permutation test (Post-R vs. all others) and Kruskal–Wallis test. (E) Same as (D) for NFKB1 regulon. (F) NF-κB - related cytokine expression (TNF, IL6, IL1B, IL1A) across 13 major cell types. Dot size, percentage of expressing cells; color, mean expression. (G) NicheNet-predicted ligand activity scores for the top 15 ligands from IL1B^+^ macrophages across receiver cell types. NF-κB -related ligands highlighted in red. (H) Radar plot comparing NF-κB pathway NES between pre- and post-treatment across cell types. (I) GSEA enrichment curves for TNF-α Signaling via NF-κB in fibroblasts and epithelial cells (post-treatment R vs. NR). (J) CD274 (PD-L1) expression in monocytes/macrophages, epithelial cells, fibroblasts, and DC cells between post-treatment R and NR. One-tailed Mann–Whitney U test. (K) Scatter plot of sample-level NF-κB pathway score versus CD8^+^ Tex exhaustion score. Spearman correlation. (L) IL-6/JAK/STAT3 pathway score in CD4^+^ T cells between post-treatment R and NR. One-tailed Mann–Whitney U test. (M) Same as (K) for Th17 score. (N) Hallmark pathway enrichment (GSEA) in monocytes/macrophages, epithelial cells, fibroblasts, and DC cells (post-treatment NR versus R). Dot size indicates significance tier; color indicates normalized enrichment score (NES). For all boxplots and scatter plots (J–M), each dot represents one sample. *, *P* < 0.05.

Within IL-1β^+^ macrophages, four cytokines—TNF, IL1B, IL6, and IL1A—were prominently expressed (**Fig. 5C**). These constitute the canonical pro-inflammatory cytokine axis: TNF-α, IL-1β, and IL-6 synergistically activate NF-κB signaling in recipient cells, while IL-1α amplifies autocrine and paracrine inflammatory loops. NF-κB serves as a master regulator linking chronic inflammation to cancer progression and therapy resistance (36,37), making the convergence on this pathway particularly relevant to understanding acquired resistance mechanisms. Gene regulatory network analysis (SCENIC) provided a transcription factor–level explanation for the inflammatory phenotype. BACH1 regulon activity was significantly reduced in IL-1β^+^ macrophages from post-treatment responders (P = 0.02, exact permutation test), whereas non-responders failed to suppress BACH1 activity (**Fig. 5D**), suggesting that persistent BACH1 regulon activity sustains the inflammatory program in resistant tumors. Similarly, NFKB1 regulon activity was elevated in non-responder IL-1β+ macrophages (P = 0.02, exact permutation test; **Fig. 5E**), consistent with NF-κB acting as a direct transcriptional driver of TNF, IL1B, IL6, and IL1A production.

Among all major cell types in the TME, monocytes/macrophages were the predominant source of these four cytokines (**Fig. 5F**). Although dendritic cells showed similar expression patterns, their substantially lower abundance in the TME suggests that monocytes/macrophages are the primary drivers of cytokine-mediated signaling in this context. NicheNet ligand–receptor analysis confirmed this: TNF was identified as the top-ranked ligand predicted to influence recipient populations, followed by IL1B, IL6, and IL1A (**Fig. 5G**)—the same four cytokines concentrated in IL-1β^+^ macrophages. Receptor expression analysis indicated that epithelial cells, fibroblasts, endothelial cells, and multiple immune subsets all express cognate receptors for these macrophage-derived cytokines, providing the molecular basis for pan-TME signal propagation.

We then examined whether this macrophage-derived signaling translates into pathway activation in recipient populations. Systematic GSEA across all major cell types revealed that every population exhibited elevated NF-κB pathway activity in non-responders (**Fig. 5H**), representing a coordinated pan-TME inflammatory state. The radar plot comparison between pre-and post-treatment samples showed that this universal NF-κB activation was specific to post-treatment non-responders, consistent with an acquired rather than pre-existing phenomenon. Individual enrichment analysis confirmed significant enrichment of TNFα/NF-κB target genes in epithelial cells from non-responders (NES = 1.75, FDR = 0.001), with a concordant trend in fibroblasts (NES = 1.47, FDR < 0.001) (**Fig. 5I**).

We then assessed the functional consequences of this pan-TME NF-κB cascade. PD-L1 expression was significantly elevated in post-treatment non-responders compared to responders across monocytes/macrophages, epithelial cells, fibroblasts, and dendritic cells (**Fig. 5J**, **Fig. S6**), and sample-level NF-κB pathway scores correlated positively with CD8^+^ T cell exhaustion (Tex; ρ = 0.74, P < 10^-6^; **Fig. 5K**), linking NF-κB-driven PD-L1 upregulation to adaptive immune dysfunction. IL-6/JAK/STAT3 pathway scores were elevated in CD4^+^ T cells from non-responders (**Fig. 5L**), and NF-κB scores correlated with Th17 signature scores (ρ = 0.40, P = 0.02; **Fig. 5M**), suggesting cytokine-driven CD4^+^ T cell reprogramming. We next examined the broader downstream consequences of this NF-κB cascade across the TME. Systematic GSEA across monocytes/macrophages, epithelial cells, fibroblasts, and dendritic cells revealed that NF-κB-driven pathways were enriched in all four cell types in post-treatment non-responders, with cell-type-specific functional manifestations (**Fig. 5N**). Epithelial cells showed enrichment for inflammatory response (NES = 1.48, P = 0.02) and hypoxia (NES = 1.56, P = 0.006), both established NF-κB-regulated programs that promote tumor cell survival and plasticity. Fibroblasts exhibited enrichment for inflammatory response (NES = 1.30, P = 0.02), consistent with NF-κB-driven stromal activation. Monocytes/macrophages themselves showed the broadest downstream activation, including inflammatory response (NES = 1.67), EMT (NES = 1.66), hypoxia (NES = 1.53), and angiogenesis (NES = 1.50; all P ≤ 0.05), reflecting both autocrine NF-κB amplification and their role as the primary source of pro-inflammatory signaling. These convergent yet cell-type-specific pathway enrichments demonstrate that macrophage-derived NF-κB signaling propagates across tumor, stromal, and immune compartments, driving coordinated TME reprogramming that extends well beyond checkpoint ligand upregulation.

Together, these results delineate a complete signaling cascade: IL-1β^+^ macrophages, driven by BACH1 and NFKB1 regulon activity, secrete TNF-α, IL-1β, IL-6, and IL-1α as the predominant cytokine source in the TME. These cytokines activate NF-κB signaling across epithelial, stromal, and immune compartments, resulting in coordinated PD-L1 upregulation, EMT, and CD8^+^ T cell exhaustion. These findings suggest that therapeutic targeting of either IL-1β^+^ macrophage recruitment/differentiation or downstream NF-κB signaling may represent strategies to overcome acquired resistance in gastric cancer.

## Discussion

Our analyses reveal two distinct and temporally separated resistance mechanisms in gastric cancer patients receiving anti-PD-1-based chemo-immunotherapy (**Fig. 6**). The first pathway involves CEACAM5/6^+^ cancer cells that are present before treatment initiation, representing intrinsic resistance. These cells establish an immunosuppressive spatial niche through macrophage recruitment and exclusion of adaptive immune cells, limiting baseline sensitivity to checkpoint blockade. The second pathway emerges during treatment, characterized by expansion of IL-1β^+^ macrophages that drive pan-TME NF-κB activation, promoting EMT, chronic inflammation, and widespread PD-L1 upregulation. CEACAM5/6 expression is detectable at baseline before macrophage expansion becomes prominent, while the macrophage-driven inflammatory cascade is most evident in post-treatment non-responders. Although the CEACAM5/6 spatial niche includes local myeloid recruitment, the NF-κB-driven pan-TME inflammatory program represents a broader, treatment-emergent phenomenon.

**Figure 6.**
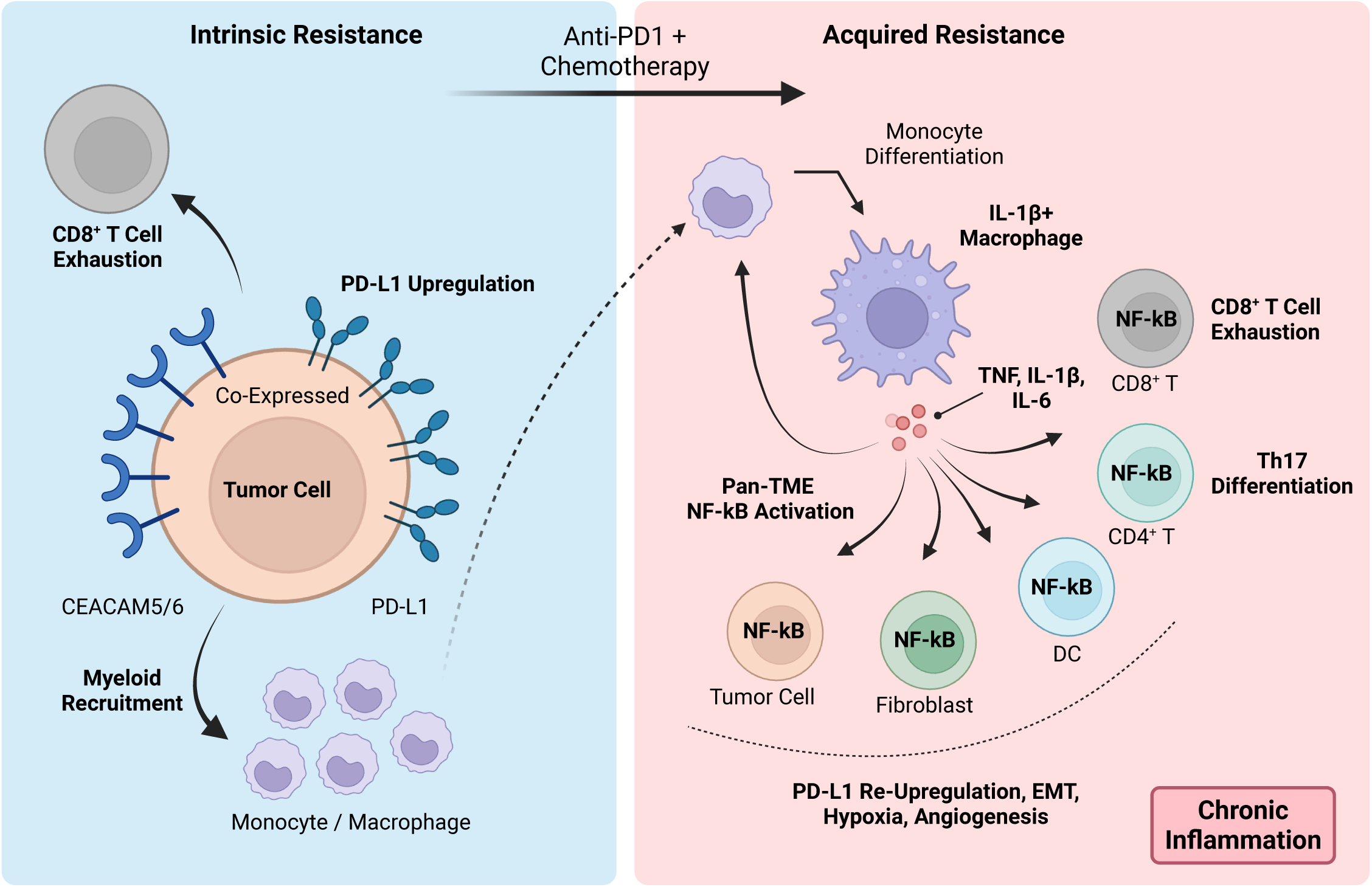
A mechanistic model of intrinsic and acquired resistance to anti-PD-1-based chemo-immunotherapy in gastric cancer. Schematic illustration. In pretreatment non-responders, CEACAM5/6^+^ tumor cells establish an immunosuppressive spatial niche through macrophage recruitment and exclusion of adaptive immune cells, limiting baseline sensitivity to immune checkpoint blockade. In posttreatment non-responders, IL1B^+^ macrophages, which activate NF-κB signaling and secrete pro-inflammatory cytokines (IL1B, TNF, IL6, IL1A). This drives upregulation of PD-L1 on tumor cells, fibroblasts, macrophages and DC; EMT in epithelial cells; angiogenesis and inflammatory responses in fibroblasts; and CD8^+^ T cell exhaustion. The CEACAM5/6–macrophage–NF-κB axis sustains immune evasion and resistance to chemoimmunotherapy.

Our findings both complement and extend recent single-cell atlases of gastric cancer (10–14). Previous studies have identified candidate resistance-associated populations—including NKG2A^+^ T cells (15), CLEVER-1^+^ macrophages (16), inflammatory CAFs, and SPP1^+^ macrophages (17,18)—but the temporal dimension of resistance has remained largely unexplored. Kumar et al. identified macrophage heterogeneity in treatment-naïve tumors but did not examine post-treatment dynamics, while Sun et al. characterized T cell exhaustion trajectories without linking them to specific tumor-intrinsic features. Our two-timepoint design reveals that these phenomena are interconnected: CEACAM5/6^+^ tumor cells are associated with spatial immune exclusion—limiting lymphocyte infiltration into the tumor core—while the CD8^+^ T cells that do infiltrate show elevated exhaustion signatures. IL-1β^+^ macrophages amplify this immunosuppressive state during treatment. The SPP1^+^ macrophages linked to immunosuppression across cancer types (17,18) may overlap with the IL-1β^+^ subset we identify, as both share inflammatory and tissue-remodeling functions, and CLEVER-1^+^ macrophages (16) may similarly represent complementary myeloid states within our framework. How NKG2A^+^ T cells relate to the exhausted CD8^+^ T cell states we observe, and how inflammatory fibroblast connect to the stromal NF-κB activation we detect, warrants further investigation.

The identification of CEACAM5/6 as a mediator of intrinsic resistance has significant implications for patient stratification and therapeutic development. CEACAM5 (CEA) has long been recognized as a tumor marker, but its functional role in immune evasion has only been appreciated recently (19,21,22). The CEACAM5/6-associated immunosuppressive niche may physically limit lymphocyte access to tumor cells, rendering PD-1 blockade ineffective regardless of T cell reinvigoration. Importantly, several CEACAM-targeting therapeutics are currently in clinical development, including antibody–drug conjugates and bispecific antibodies (23,40–42). Our results support the use of CEACAM5/6 as a biomarker for patient selection for standard anti-PD-1–based chemo-immunotherapy and further suggest that combining these agents with anti-PD-1 therapy may benefit patients with high baseline CEACAM5/6 expression.

The IL-1β^+^ macrophage-driven NF-κB cascade we identified as a hallmark of acquired resistance aligns with emerging evidence implicating tumor-associated macrophages as central regulators of immunotherapy response (17,18,32–34). The observation that IL-1β^+^ macrophages are markedly reduced in responders but expanded in non-responders suggests that this population may represent a therapeutic target. IL-1β has been linked to tumor-promoting inflammation and immunosuppression across cancer types, and the CANTOS trial demonstrated that IL-1β inhibition with canakinumab reduced cancer incidence and mortality in atherosclerosis patients (43). Our finding that macrophage-derived IL-1β drives downstream IL-6 production, EMT, and PD-L1 upregulation (44–46) provides a mechanistic rationale for combining IL-1β or NF-κB pathway inhibitors with checkpoint blockade. Several clinical trials exploring such combinations are underway, and our biomarker framework may help identify patients most likely to benefit.

Together, our study suggests that patients may benefit from different therapeutic strategies depending on which resistance pathway predominates. Rather than uniform combination regimens, our data argue for sequential strategies tailored to the dominant resistance mechanism: CEACAM-targeting agents at treatment initiation for patients with high baseline CEACAM5/6, followed by anti-inflammatory intervention upon disease progression. Prospective clinical trials incorporating temporal biomarker assessment are warranted to translate this framework into clinical benefit.

## Acknowledgments

This work was supported by the National Natural Science Foundation of China (Grant Nos. 92474106, 82573742, and 82403738), the Noncommunicable Chronic Diseases–National Science and Technology Major Project (Grant No. 2024ZD0520600), and the “Leading Goose” R&D Program of Zhejiang (Grant No. 2026C02A1103); and by the Barnhart Family Distinguished Professorship in Targeted Therapies at The University of Texas MD Anderson Cancer Center.

## Methods

### Study Design and Patient Cohort

Patients with gastric cancer receiving anti-PD-1-based chemo-immunotherapy (anti-PD-1: Nivolumab/ Tislelizumab/Sintilimab; chemo: mFOLFOX6/SOX) were enrolled at the Second Affiliated Hospital, Zhejiang University School of Medicine between 2021 and 2023. Exclusion criteria included presence of other concurrent malignancies; active or history of autoimmune diseases requiring systemic treatment; inadequate organ function or severe systemic diseases; regular use of systemic corticosteroids or other immunosuppressants prior to treatment. Treatment response was evaluated according to RECIST v1.1 criteria, with responders (R) defined as patients achieving complete or partial response (CR/PR) and non-responders (NR) defined as those with stable or progressive disease (SD/PD). Pre-treatment samples were collected prior to chemo-immunotherapy initiation, and post-treatment samples were obtained after therapy. A total of 35 patients contributed 70 samples across multiple tissue sites, including stomach (n = 32), liver (n = 12), peripheral blood (n = 11), lymph node (n = 10), ovary (n = 3), and metastatic lymph node (n = 2). For the primary response analysis in stomach, 8 pre-treatment samples (4 R, 4 NR) and 11 post-treatment samples (5 R, 6 NR) with confirmed response classifications following neoadjuvant chemo-immunotherapy were used. This study was approved by the Institutional Review Board of the Second Affiliated Hospital of Zhejiang University (Approval protocol number: I20221149). Written informed consent was obtained from all patients.

### Sample Collection and Processing

Fresh tumor tissues were stored in the sCelLive Tissue Preservation Solution (Singleron Biotechnologies) on ice within 30 min after surgery or biopsy. Specimens were washed with Hanks Balanced Salt Solution three times and minced into 1-2 mm pieces. Tissues were digested with GEXSCOPE Tissue Dissociation Mix (Singleron Biotechnologies) at 37℃ for 15 min with continuous agitation. Following digestion, samples were filtered through 40-um sterile strainers and centrifuged at 1,000 rpm for 5 min. Cell pellets were resuspended and incubated with GEXSCOPE Red Blood Cell Lysis Buffer (Singleron Biotechnologies) for 10 min at room temperature to remove erythrocytes. After washing with PBS, cell viability was assessed using trypan blue staining, and samples with >80% viability proceeded for single-cell library preparation.

### Single-cell RNA Sequencing

Single-cell suspensions (1 × 10^5^ cells/mL) were loaded onto Singleron microfluidic devices. The scRNA-seq libraries were constructed using the GEXSCOPE Single Cell RNA Library Kit (Singleron Biotechnologies). Individual cells were captured in droplets containing barcoded beads, followed by cell lysis, mRNA capture, reverse transcription, and cDNA amplification. Libraries were sequenced on the Illumina HiSeq X10 platform with 150 bp paired-end reads.

### Data Processing and Quality Control

Raw sequencing reads were processed using the CeleScope pipeline (Singleron Biotechnologies) for read alignment to the human reference genome (GRCh38) and gene expression quantification. Quality control was performed in three phases. In the first phase (droplet-level filtering), overall data quality was assessed and empty droplets were removed using EmptyDrops from the DropletUtils R package. In the second phase (cluster-informed filtering), Scrublet was used for doublet detection, and cells were further filtered based on clustering information to identify collectively abnormal cell populations. In the third phase (biology-informed filtering), prior biological knowledge was leveraged to identify cluster-specific markers and remove cells with inconsistent expression patterns. Standard quality filters were applied: cells with fewer than 200 detected genes or fewer than 500 UMI counts were excluded.

### Data Integration and Normalization

Gene expression counts were normalized using log-normalization with a target sum of 10,000 UMI per cell. Highly variable genes were identified using scanpy (47) (min_mean = 0.0125, max_mean = 3, min_disp = 0.5). Total counts and mitochondrial percentage were regressed out, and gene expression values were scaled with a maximum value of 10. Principal component analysis (50 components) was performed, followed by Harmony batch correction (48) across samples (max_iter_harmony = 20). Neighborhood graphs were constructed using 10 neighbors and 50 principal components, and UMAP was used for two-dimensional visualization. Leiden clustering (resolution = 0.3) was applied for unsupervised cluster identification. Sub-clustering of specific cell populations was performed using the same pipeline at higher resolutions as needed. All analyses were performed using Python 3.10 with scanpy 1.10.4, anndata 0.11.1, scipy 1.14.1, matplotlib 3.9.3, and seaborn 0.13.2. Gene regulatory network analysis used pySCENIC 0.12.1, and pathway enrichment analysis used GSEApy 1.1.4. Cell type deconvolution was performed using BayesPrism v2.2.2 in R.

### Cell Type Annotation

Unsupervised clustering was performed using the Leiden algorithm (49). A total of 14 major cell types were identified: B cells, CD4^+^ T cells, CD8^+^ T cells, dendritic cells, endothelial cells, epithelial cells, fibroblasts, hepatocytes, mast cells, monocytes/macrophages, NK cells, neutrophils, pericytes, and plasma cells. Of these 14 cell types, hepatocytes and pericytes were identified only in non-stomach tissues; stomach-specific analyses utilized 12 cell types (excluding hepatocytes and pericytes). NicheNet ligand-receptor analysis included 13 receiver cell types (12 stomach types plus pericytes, which were present in the stomach pre-treatment responder subset). Major cell types were annotated based on canonical marker gene expression. Wilcoxon rank-sum test was used for cluster marker identification during annotation. Sub-clustering identified 84 minor cell states.

### Epithelial Metaprogram Discovery

Gene expression programs in malignant epithelial cells were identified using per-sample non-negative matrix factorization (NMF). For each sample with ≥100 epithelial cells, expression data were normalized to counts per million (CPM) and log2-transformed. NMF was performed using the Brunet multiplicative update algorithm (R NMF package) across K values of 4 through 9, with 10 independent runs per K value (random seed = 42). The top 50 genes by weight were extracted from each program. Programs were filtered through a three-step robust filtering process. First, within-sample recurrence required each program to have ≥1 matching program (from a different K value in the same sample) sharing ≥35 of 50 top genes. Second, cross-sample recurrence required each program to have ≥1 matching program from any other sample sharing ≥10 of 50 top genes. Third, redundancy removal discarded programs overlapping >10 genes with a higher-ranked program within the same sample. Filtered programs were clustered into consensus metaprograms using Jaccard similarity. Programs with Jaccard index ≥0.10 were grouped via greedy seed-expansion (minimum 3 programs per metaprogram). Consensus gene signatures were defined by ranking genes by frequency across member programs (descending), then by average NMF weight (descending), and selecting the top 50 genes. Metaprogram activity scores per cell were calculated using scanpy score_genes (ctrl_size = 50, n_bins = 25), and aggregated to sample-level means.

### Differential Abundance Analysis

Differential abundance analysis was performed using Milo (51) (pertpy Python implementation) to compare cell type proportions between responders and non-responders. Neighborhoods were constructed with K = 15 neighbors and proportion = 0.1. Differential abundance was tested using the edgeR framework with significance defined as SpatialFDR < 0.1 (52). When the cell population exceeded 8,000 cells, random subsampling was applied prior to neighborhood construction.

### Differential Expression Analysis

Differential expression analysis between responders and non-responders was performed using the MAST hurdle model (53). The model was specified as: zlm(∼ condition + sample_id + cngeneson), where condition encodes response status (reference: non-responder), sample_id accounts for patient-level effects, and cngeneson (scaled cellular detection rate) controls for technical variation. Genes expressed in <10% of cells were excluded. P-values were adjusted using the Benjamini-Hochberg method, with adjusted P < 0.05 considered significant.

### Gene Set Enrichment Analysis

Gene set enrichment analysis (GSEA) was performed using GSEApy (prerank method). Genes were ranked by log_2_FC × -log_10_(P-value) from MAST results. Enrichment was assessed against MSigDB Hallmark 2020 gene sets with 1,000 permutations, minimum gene set size of 15, and maximum of 500. NF-κB pathway activation was assessed using the Hallmark TNFα Signaling via NF-κB gene set. EMT scores were calculated using the Hallmark EMT gene set. Pathway activity scores per cell were calculated using scanpy score_genes. All gene signatures used in this study are listed in Table S3.

### Cell-Type-Specific NF-κB Pathway Enrichment

For NF-κB pathway enrichment analysis across cell types (Fig. 5H, I), differentially expressed genes were identified between non-responders and responders using Welch’s t-test with overestimated variance (scanpy.tl.rank_genes_groups, method = ’t-test_overestim_var’), applied separately to pre-treatment and post-treatment stomach samples. For cell populations exceeding 5,000 cells, random subsampling was performed (n = 5,000; random_state = 42). Genes were ranked by the product of log_2_ fold change and −log_10_(P-value). GSEA was then performed using GSEApy prerank against MSigDB Hallmark 2020 gene sets (min_size = 5, max_size = 500; 1,000 permutations; seed = 42). Normalized enrichment scores (NES) for the TNFα Signaling via NF-κB pathway were extracted for each of the 13 major cell types.

### Regulon Analysis

Gene regulatory network analysis was performed using pySCENIC (54) on monocytes/macrophages (68,289 cells). The pipeline consisted of three steps: (i) gene regulatory network inference using GRNBoost2 with human transcription factors, (ii) regulon identification via cisTarget motif enrichment using motifs-v10nr_clust-nr.hgnc-m0.001-o0.0.tbl annotations and feather-format ranking databases, and (iii) regulon activity scoring using AUCell. Regulon activity was quantified as AUCell scores per cell.

### Cell-Cell Communication Analysis

Ligand-receptor interaction analysis was performed using NicheNet (55) to identify signaling from the inflammatory macrophage subset (IL-1β^+^ inflammatory macrophage subset, sender) to 13 receiver cell types. Analysis was restricted to post-treatment samples, comparing non-responders versus responders, to identify signaling pathways associated with acquired resistance. Ligands expressed in >10% of sender cells were considered. For each receiver, the top 250 most variable genes among NicheNet target genes were selected, and ligand activity was scored using Pearson correlation. The top 30 ligands per receiver were reported.

### Immune Module Identification

To identify co-regulated immune cell modules, we employed an ecotype-inspired approach. Sample-level proportions of each minor immune cell state were calculated within stomach immune cells using within-immune normalization. Spearman rank correlation coefficients were computed across all cell state proportion vectors. Hierarchical clustering was performed on the correlation matrix using Ward’s linkage method. The dendrogram was cut at K = 5 to define five immune modules. Module proportions per sample were calculated as the mean proportion of constituent cell states, followed by row normalization. The five modules comprised: M1, T/NK/DC (20 cell states); M2, monocyte/macrophage (6 cell states including IL-1β^+^ inflammatory macrophages); M3, mixed immune (14 cell states including Tfh, effector T, and activated DC); M4, neutrophil (8 cell states); and M5, B/plasma (8 cell states).

### Immunohistochemistry

CEACAM5 and CEACAM6 immunohistochemical staining was performed on formalin-fixed paraffin-embedded tissue sections from 8 patients (4 responders and 4 non-responders, Table S5). Primary antibodies used were anti-CEACAM5 (clone EPCEAR7, Abcam, catalog ab133633; dilution 1:2000) and anti-CEACAM6 (clone EPR4403, Abcam, catalog ab134074; dilution 1:200). Staining was performed using a manual protocol. Heat-induced antigen retrieval was performed using EDTA buffer (pH 9.0) in a pressure cooker for 15 min. Stained slides were digitized using a KFBIO KF-PRO-120 digital pathology slide scanner (Konfotech, Ningbo, China) at 40× magnification (0.25 μm/pixel resolution). IHC images were quantified using Hematoxylin-DAB color deconvolution (58) implemented in scikit-image. The DAB optical density channel was thresholded (OD > 0.02) within tissue regions to identify positively stained areas. Staining area percentage was calculated as DAB-positive pixels divided by total tissue pixels. Combined CEACAM5 + CEACAM6 staining area was calculated as the sum of CEACAM5 and CEACAM6 positive staining percentages per patient. Statistical comparison between responder and non-responder groups was performed using one-tailed Mann–Whitney U test.

### TCGA-STAD Bulk RNA-seq Validation

TCGA-STAD RNA-seq data (FPKM) were downloaded from the GDC Data Portal (50). The original dataset contained 410 sample columns. After quality assessment, we identified and removed 3 normal-only samples and corrected 14 samples where normal tissue expression had been used in place of tumor expression, yielding a cleaned matrix of 407 tumor-only samples. Ensembl gene IDs were converted to gene symbols, retaining the highest-expressed gene for duplicates. Clinical data including vital status, days to death, and days to last follow-up were obtained for survival analyses. Cell type deconvolution of TCGA-STAD bulk RNA-seq data was performed using BayesPrism (v2.2.2) (56) in R. The single-cell reference consisted of 6,834 cells across 5,020 genes from our scRNA-seq dataset. Bulk expression input was provided as gene expression profiles. Outlier genes were filtered (outlier.cut = 0.01, outlier.fraction = 0.1). Cell-type-specific expression profiles and cell type fractions were estimated for 407 tumor-only TCGA-STAD samples.

### PRJEB25780 (TIGER) Bulk RNA-seq Validation

Bulk RNA-seq data from the PRJEB25780 (TIGER) cohort of gastric cancer patients treated with anti-PD-1 therapy were collected. The cohort comprised 45 treated tumor samples (12 responders, 33 non-responders). Cell type deconvolution was performed using BayesPrism (v2.2.2) with the same single-cell reference (6,834 cells × 5,020 genes) and parameters as described above. Epithelial-specific deconvolved expression profiles were extracted for CEACAM gene analysis. Details of all external cohorts are provided in Table S2.

### Tumor Purity Estimation

Tumor purity of PRJEB25780 samples was estimated using the ESTIMATE algorithm (v1.0.13) with the Illumina platform setting. Stromal, immune, and ESTIMATE scores were computed per sample, and tumor purity was derived using the cosine transformation formula. CEACAM expression comparisons between responders and non-responders were adjusted for tumor purity using linear models (expression ∼ response + TumorPurity).

### Spatial Transcriptomics Analysis

Spatial transcriptomics data from an independent Korean Gut Atlas cohort (GEO: GSE251950; 10 Visium samples) were analyzed for external validation. Cell type deconvolution was performed using GraphST (57) (epochs = 1,200, learning_rate = 0.001) with spatial graphs constructed using 6 nearest neighbors based on hexagonal Visium geometry. The single-cell reference comprised 14 cell types: 12 non-epithelial cell types plus CEACAM-high and CEACAM-low epithelial populations, split based on *CEACAM5* and *CEACAM6* expression. Cell-to-spot projection used retain_percent = 0.15. Spatial domains were identified by Leiden clustering (resolution = 0.5) on GraphST embeddings. To assess spatial immune exclusion by CEACAM-expressing tumor cells, linear mixed-effects models were fitted: Immune ∼ CEACAM_ratio + Total_Epi + (1|Sample), where CEACAM_ratio represents the proportion of CEACAM-high epithelial cells among total epithelial cells per spot, and Total_Epi controls for overall tumor cell density. Models were fitted using statsmodels (LBFGS optimizer, maxiter = 1,000) on spots containing at least 5% epithelial content.

### External Single-Cell RNA-seq Validation

Two independent gastric cancer scRNA-seq datasets (GSE183904, GSE239676) were used for external validation of the IL-1β^+^ macrophages inflammatory macrophage signature. Both datasets were annotated using CellTypist (Immune_All_Low.pkl model, majority_voting = True). IL-1β+ macrophage signature scores were computed using scanpy score_genes with a 15-gene inflammatory macrophage signature (*IL1B, TNF, CXCL8, CCL3, CCL4, CXCL2, CXCL3, IL1A, SOD2, NFKBIA, PTGS2, G0S2, IER3, PLAUR, EREG*) derived from the top markers of the IL-1β^+^ inflammatory macrophage cluster in our primary dataset. IL-1β^+^ macrophage proportions per sample were compared between tumor and normal tissue groups using the Mann-Whitney U test.

### Patient Survival Analysis

Kaplan-Meier survival analysis was performed on TCGA-STAD patients. For *CEACAM5* and *CEACAM6* expression, BayesPrism-deconvolved epithelial expression values were log_2_-transformed and patients were stratified by median split into high and low groups. Survival differences were assessed using the log-rank test (Python lifelines package). For macrophage subset proportion analysis, optimal expression cutpoints were determined using surv_cutpoint (R survminer package, MINPROP = 0.3), and Cox proportional hazards regression was used to estimate hazard ratios with 95% confidence intervals.

### Bootstrap Confidence Intervals

Confidence intervals for fold changes in monocyte/macrophage cell state proportions between responders and non-responders were estimated using the percentile bootstrap method. Sample-level proportions were resampled with replacement (10,000 iterations, seed = 42) separately within responder (n = 5) and non-responder (n = 6) groups. For each iteration, the fold change was calculated as the ratio of resampled group means (R/NR). The 95% confidence interval was defined by the 2.5th and 97.5th percentiles of the bootstrap distribution.

## Statistical Analysis

Statistical analyses were performed using Python (v3.10) and R (v4.2). Two-group comparisons were performed using the Mann-Whitney U test. One-tailed tests were used for directional hypotheses where non-responder values were hypothesized to exceed responder values (e.g., *CEACAM5*/6 expression, *PD-L1 expression*, EMT scores). The epithelial subpopulation proportion comparison (Fig. 2f) was evaluated with a two-tailed test given the exploratory nature of this comparison. Paired comparisons in spatial data were assessed using the Wilcoxon signed-rank test. Multi-group comparisons used the Kruskal-Wallis test. Correlations were assessed using Spearman rank correlation. For small-sample comparisons (n = 4 vs. n = 4), exact permutation tests using all possible permutations were performed to obtain precise P-values. Differential expression analysis was performed using the MAST hurdle model as described above. All comparisons were performed at the sample level using sample means, rather than at the cell level, to account for the non-independence of cells originating from the same patient, unless otherwise specified. P-values < 0.05 were considered statistically significant. One sample (GC_1228) was identified as a statistical outlier for PD-L1 expression (Grubbs test, G = 5.43, P < 0.05) and excluded from the CEACAM–PD-L1 correlation analysis (Fig. 3a).

## Data Availability

Processed single-cell RNA sequencing data and H&E/IHC staining generated in this study will be available upon publication. Spatial transcriptomic data used for validation are publicly available from GEO under accession number GSE251950. Two independent gastric cancer scRNA-seq datasets used for external validation are available from GEO under accession numbers GSE239676 and GSE183904. Bulk RNA-seq data from the TIGER immunotherapy cohort are available from the European Nucleotide Archive under accession number PRJEB25780. TCGA-STAD data were obtained from the NCI GDC Data Portal.

## Supplementary Figures

**Figure S1.** Quality control, batch integration, and cell type annotation of scRNA-seq data

**Figure S2.** CEACAM5/6 validation and meta-program characterization

**Figure S3.** CD8^+^ T cell sub-cluster characterization and exhaustion markers

**Figure S4.** Spatial transcriptomic validation across all GSE251950 samples

**Figure S5.** Immune module proportions across treatment–response groups

**Figure S6.** CD274 (PD-L1) expression across additional cell types in post-treatment samples

## Supplementary Tables

**Supplementary Table 1.** Patient and sample characteristics.

**Supplementary Table 2.** IHC validation cohort characteristics.

**Supplementary Table 3.** External cohort details.

**Supplementary Table 4.** Gene signatures. Complete gene lists for T cell exhaustion, EMT (mesenchymal and epithelial), MDSC, and other gene signatures used for module scoring and pathway analyses.

**Supplementary Table 5.** Checkpoint and immunomodulatory receptors. Comprehensive list of immune checkpoint and immunomodulatory receptor genes analyzed by pseudo-bulk differential expression, including protein names, functional categories, representative therapeutic agents, and clinical trial identifiers.

## Notes

### Competing Interest Statement

The authors have declared no competing interest.

