## Supplementary Figures for "CEACAM5/6^+^ Tumor Cells and IL-1β^+^ Macrophages Drive Resistance to Chemo-immunotherapy in Gastric Cancer"

### Figure S1

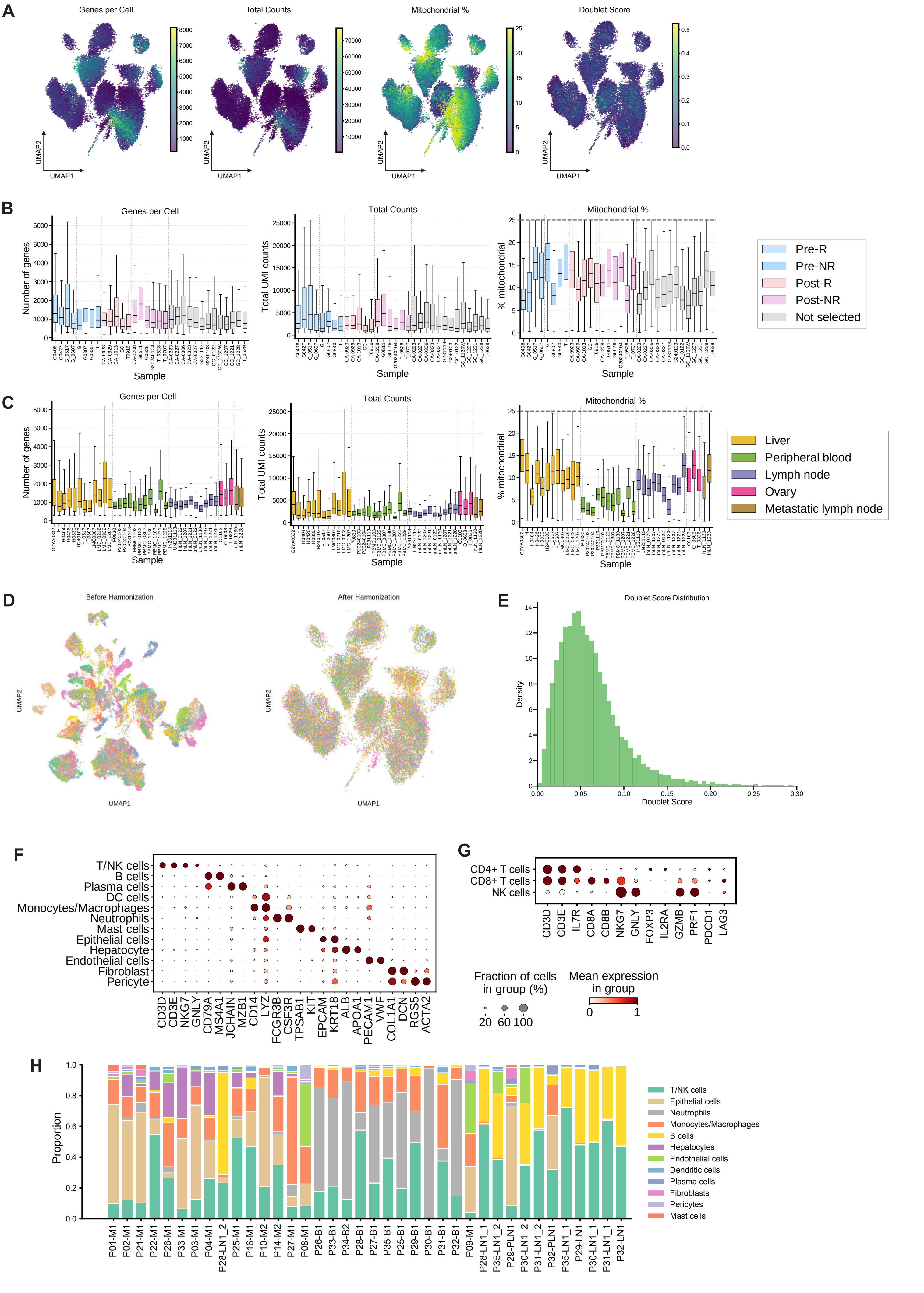
A
Genes per Cell
Total Counts
Mitochondrial %
Doublet Score
UMAP1
UMAP2
B
Genes per Cell
Number of genes
Total Counts
Total UMI counts
Mitochondrial %
% mitochondrial
Sample
Pre-R
Pre-NR
Post-R
Post-NR
Not selected
C
Liver
Peripheral blood
Lymph node
Ovary
Metastatic lymph node
D
Before Harmonization
After Harmonization
E
Doublet Score Distribution
Density
Doublet Score
F
T/NK cells
B cells
Plasma cells
DC cells
Monocytes/Macrophages
Neutrophils
Mast cells
Epithelial cells
Hepatocyte
Endothelial cells
Fibroblast
Pericyte
CD3D
CD3E
NKG7
GNLY
CD79A
MS4A1
JCHAIN
MZB1
CD14
LYZ
FCGR3B
CSF3R
TPSAB1
KIT
EPCAM
KRT18
ALB
APOA1
PECAM1
VWF
COL1A1
DCN
RGS5
ACTA2
G
CD4+ T cells
CD8+ T cells
NK cells
CD3D
CD3E
IL7R
CD8A
CD8B
NKG7
GNLY
FOXP3
IL2RA
GZMB
PRF1
PDCD1
LAG3
Fraction of cells in group (%)
20 60 100
Mean expression in group
0 1
H
Proportion
T/NK cells
Epithelial cells
Neutrophils
Monocytes/Macrophages
B cells
Hepatocytes
Endothelial cells
Dendritic cells
Plasma cells
Fibroblasts
Pericytes
Mast cells

**Figure S1. Quality control, batch integration, and cell type annotation of scRNA-seq data**
(A) UMAP embeddings colored by number of genes detected, total UMI counts, mitochondrial gene percentage, and doublet score. (B) Distribution of QC metrics across 32 stomach samples, colored by treatment–response group. (C) Distribution of QC metrics across 38 non-stomach samples, colored by tissue site. (D) UMAP colored by sample identity before (left) and after (right) Harmony batch correction. (E) Distribution of Scrublet doublet scores with predicted doublets overlaid. (F) Dotplot of canonical marker genes across 12 major cell types (T/NK cells merged). (G) Dotplot of canonical marker genes for T/NK cell subtypes (CD4+ T, CD8+ T, NK cells). (H) Cell-type composition of non-stomach samples (n = 38), displayed as stacked bar chart. Color mapping matches Figure 1C.

Figure S2

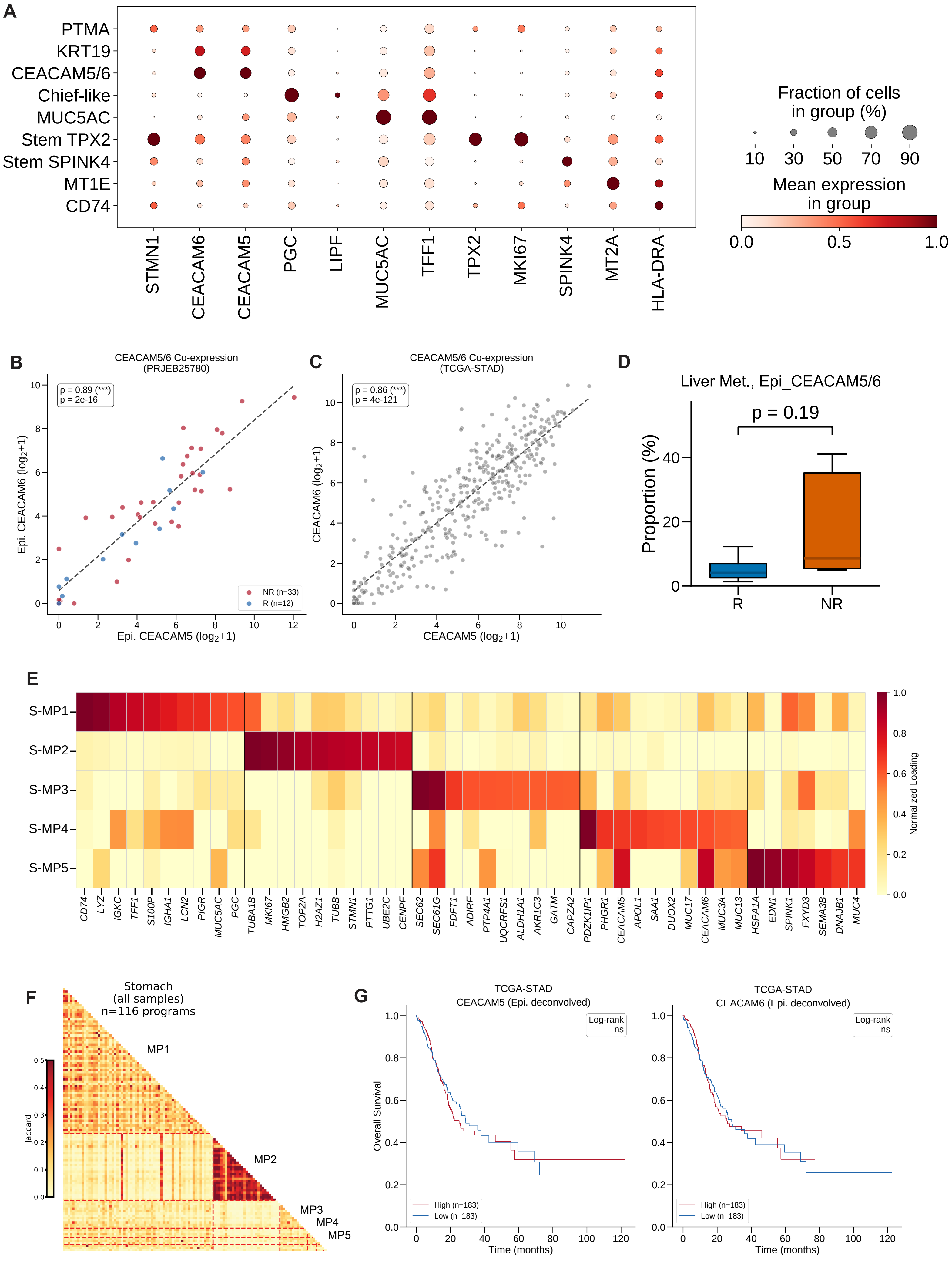

A
PTMA
KRT19
CEACAM5/6
Chief-like
MUC5AC
Stem TPX2
Stem SPINK4
MT1E
CD74
STMN1
CEACAM6
CEACAM5
PGC
LIPF
MUC5AC
TFF1
TPX2
MKI67
SPINK4
MT2A
HLA-DRA
Fraction of cells
in group (%)
10
30
50
70
90
Mean expression
in group
0.0
0.5
1.0
B
CEACAM5/6 Co-expression
(PRJEB25780)
ρ = 0.89 (***)
p = 2e-16
Epi. CEACAM6 (log2+1)
Epi. CEACAM5 (log2+1)
NR (n=33)
R (n=12)
C
CEACAM5/6 Co-expression
(TCGA-STAD)
ρ = 0.86 (***)
p = 4e-121
CEACAM6 (log2+1)
CEACAM5 (log2+1)
D
Liver Met., Epi_CEACAM5/6
p = 0.19
Proportion (%)
R
NR
E
S-MP1
S-MP2
S-MP3
S-MP4
S-MP5
CD74
LYZ
IGKC
TFF1
S100P
IGHA1
LCN2
PIGR
MUC5AC
PGC
TUBA1B
MKI67
HMGB2
TOP2A
H2AZ1
TUBB
STMN1
PTTG1
UBE2C
CENPF
SEC62
SEC61G
FDFT1
ADIRF
PTP4A1
UQCRFS1
ALDH1A1
AKR1C3
GATM
CAPZA2
PDZK1IP1
PHGR1
CEACAM5
APOL1
SAA1
DUOX2
MUC17
CEACAM6
MUC3A
MUC13
HSPA1A
EDN1
SPINK1
FXYD3
SEMA3B
DNAJB1
MUC4
Normalized Loading
F
Stomach
(all samples)
n=116 programs
MP1
MP2
MP3
MP4
MP5
Jaccard
G
TCGA-STAD
CEACAM5 (Epi. deconvolved)
Log-rank
ns
Overall Survival
Time (months)
High (n=183)
Low (n=183)
TCGA-STAD
CEACAM6 (Epi. deconvolved)
Log-rank
ns
Time (months)
High (n=183)
Low (n=183)

**Figure S2. CEACAM5/6 validation and meta-program characterization**

(A) Dotplot of marker genes across 9 epithelial sub-clusters. (B) Spearman correlation between CEACAM5 and CEACAM6 in an independent cohort (PRJEB25780; BayesPrism-deconvolved epithelial expression), colored by R versus NR. (C) Spearman correlation between CEACAM5 and CEACAM6 in TCGA-STAD (n = 407; BayesPrism-deconvolved epithelial expression). (D) Proportion of CEACAM5/6-high (Epi_CEACAM5/6) epithelial cells in liver metastasis samples, R versus NR. Mann–Whitney U test. (E) Gene signature heatmap of 5 NMF meta-programs (MP1–MP5), showing top 10 genes per program. (F) Jaccard similarity heatmap of NMF meta-programs (MP1–MP5) across all gastric tumor samples. (G) Kaplan–Meier survival curves for CEACAM5/6-high versus -low groups in TCGA-STAD (n = 407; BayesPrism-deconvolved epithelial expression, median split). Log-rank test.

Figure S3

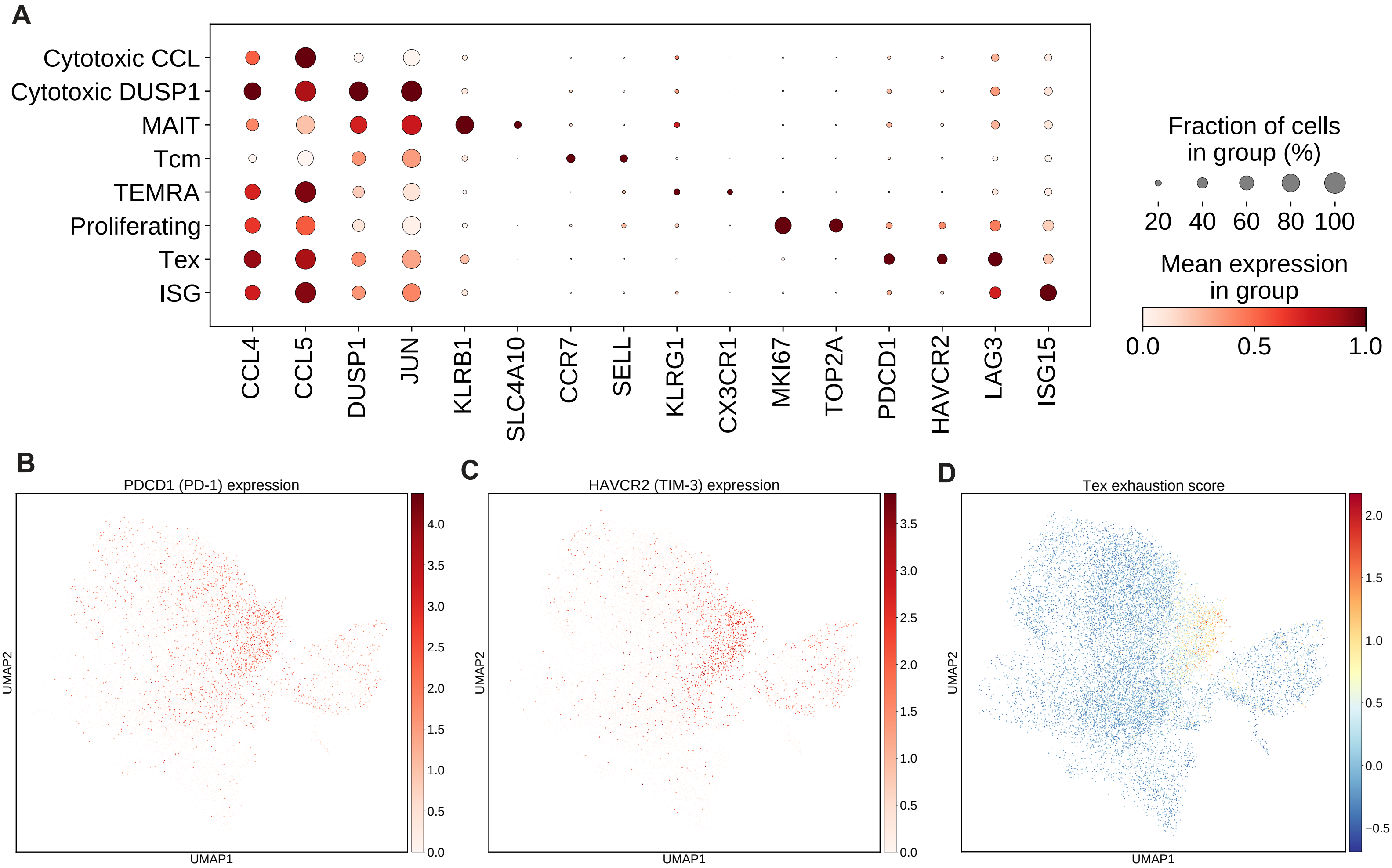


**Figure S3. CD8+ T cell sub-cluster characterization and exhaustion markers**

(A) Dotplot of marker genes across 8 CD8+ T cell sub-clusters. Dot size, percentage of expressing cells; color, mean expression. (B) UMAP of CD8+ T cells colored by PDCD1 (PD-1) expression (log-normalized). (C) UMAP of CD8+ T cells colored by HAVCR2 (TIM-3) expression (log-normalized). (D) UMAP of CD8+ T cells colored by exhaustion (Tex) score, calculated using a 10-gene signature (PDCD1, HAVCR2, LAG3, TIGIT, CTLA4, TOX, ENTPD1, LAYN, CXCL13, BATF; see Methods).

Figure S4

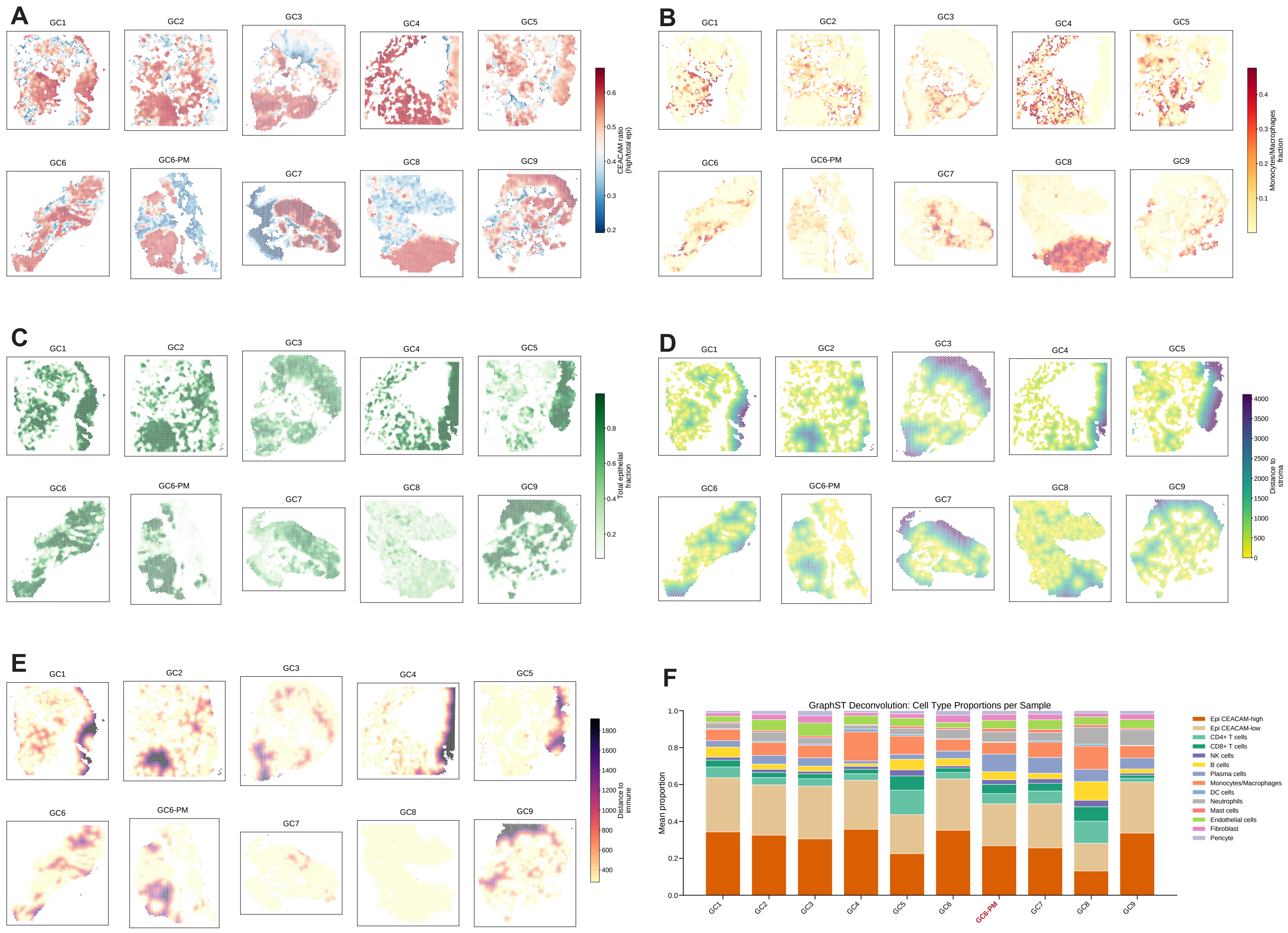


**Figure S4. Spatial transcriptomic validation across all GSE251950 samples**

(A–E) Spatial maps across all 10 gastric cancer samples showing: (A) CEACAM expression ratio (CEACAM5 + CEACAM6 deconvolved expression), (B) deconvolved monocyte/macrophage fraction, (C) epithelial neighborhood density, (D) distance to nearest stromal region, and (E) distance to nearest immune region. (F) Stacked bar chart of deconvolved cell type proportions per sample (GraphST). Color mapping matches Figure 1C.

Figure S5

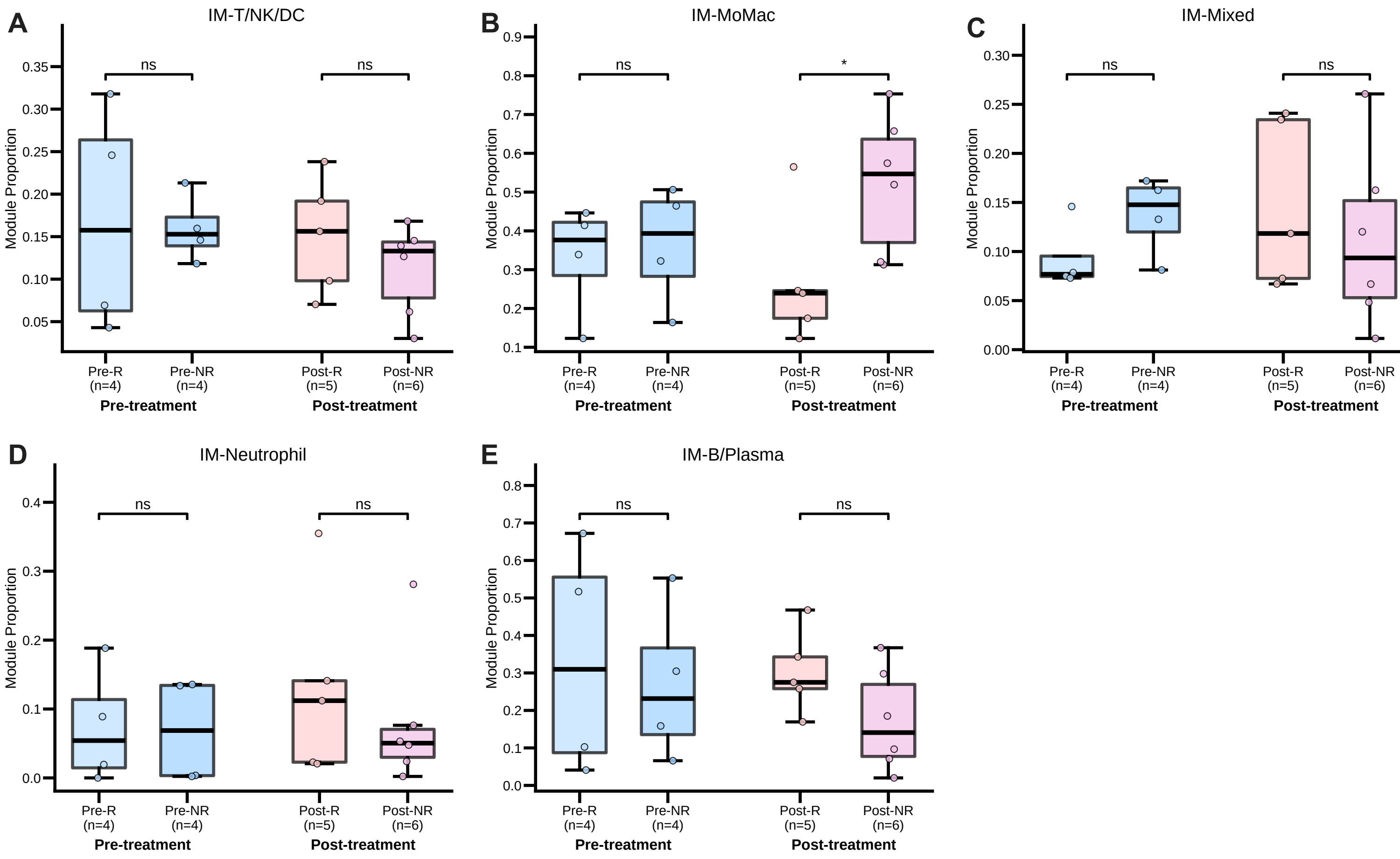


**Figure S5. Immune module proportions across treatment–response groups**

(A–E) Immune module proportions in pre-treatment and post-treatment stomach samples. Left, pre-treatment R (n = 4) versus NR (n = 4); right, post-treatment R (n = 5) versus NR (n = 6). Module 1 (IM-T/NK/DC) (A), Module 2 (IM-MoMac) (B), Module 3 (IM-Mixed) (C), Module 4 (IM-Neutrophil) (D), Module 5 (IM-B/Plasma) (E). Mann–Whitney U test. Each dot represents one sample.

Figure S6

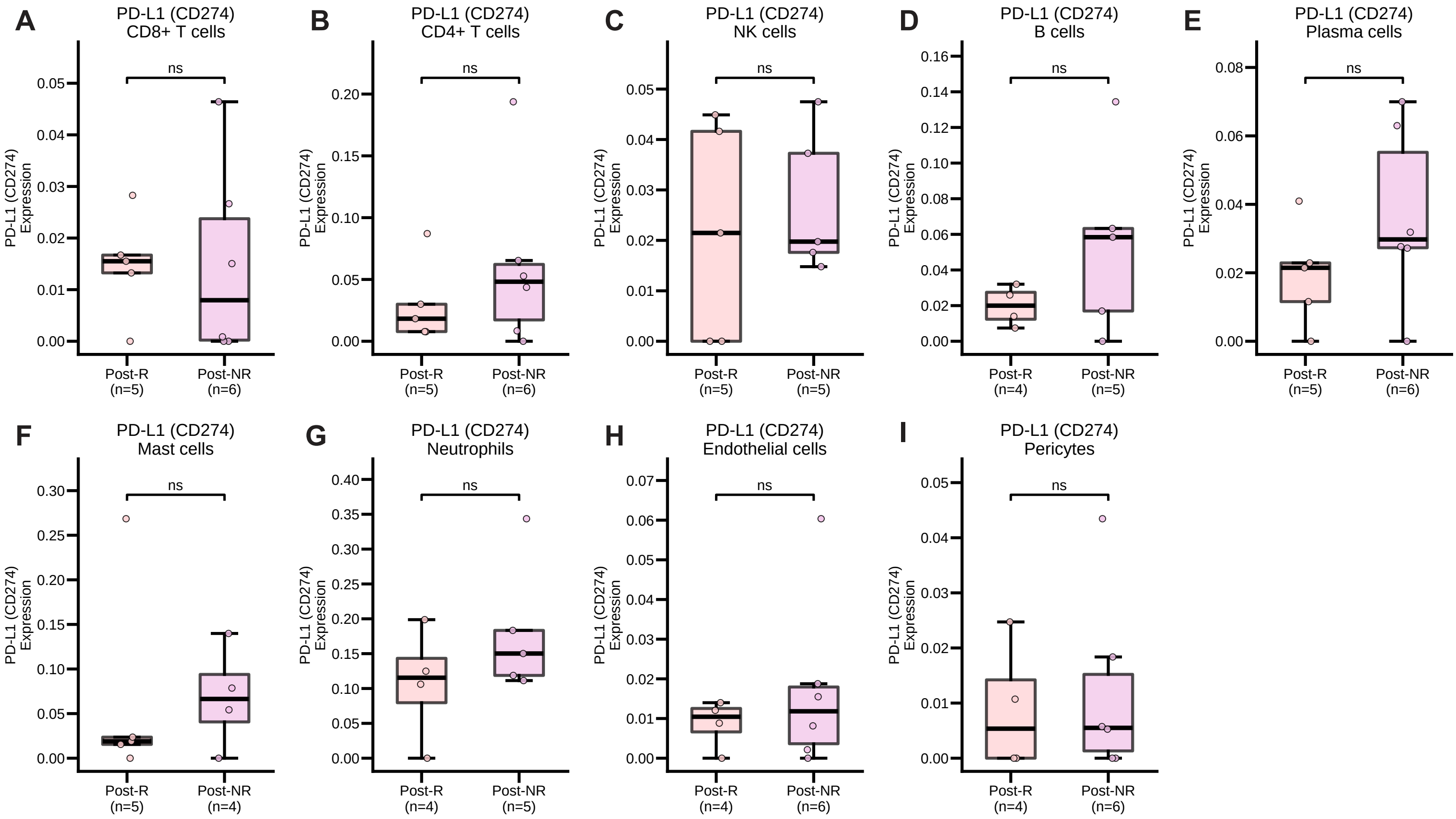


**Figure S6. CD274 (PD-L1) expression across additional cell types in post-treatment samples**
(A–I) Sample-level CD274 (PD-L1) expression in post-treatment R (n = 5) versus NR (n = 6) for cell types not shown in Figure 5: CD8+ T cells (A), CD4+ T cells (B), NK cells (C), B cells (D), Plasma cells (E), Mast cells (F), Neutrophils (G), Endothelial cells (H), and Pericytes (I). Samples with fewer than 20 cells excluded. Mann–Whitney U test. Each dot represents one sample.
